# Spatial population genomics of a recent mosquito invasion

**DOI:** 10.1101/2020.11.30.405191

**Authors:** Thomas L Schmidt, T. Swan, Jessica Chung, Stephan Karl, Samuel Demok, Qiong Yang, Matt A Field, Mutizwa Odwell Muzari, Gerhard Ehlers, Mathew Brugh, Rodney Bellwood, Peter Horne, Thomas R Burkot, Scott Ritchie, Ary A Hoffmann

## Abstract

Population genomic approaches can characterise dispersal across a single generation through to many generations in the past, bridging the gap between individual movement and intergenerational gene flow. These approaches are particularly useful when investigating dispersal in recently altered systems, where they provide a way of inferring long-distance dispersal between newly established populations and their interactions with existing populations. Human-mediated biological invasions represent such altered systems which can be investigated with appropriate study designs and analyses. Here we apply temporally-restricted sampling and a range of population genomic approaches to investigate dispersal in a 2004 invasion of *Aedes albopictus* (the Asian tiger mosquito) in the Torres Strait Islands (TSI) of Australia. We sampled mosquitoes from 13 TSI villages simultaneously and genotyped 373 mosquitoes at genome-wide single nucleotide polymorphisms (SNPs): 331 from the TSI, 36 from Papua New Guinea (PNG), and 4 incursive mosquitoes detected in uninvaded regions. Within villages, spatial genetic structure varied substantially but overall displayed isolation by distance and a neighbourhood size of 232–577. Close kin dyads revealed recent movement between islands 31–203 km apart, and deep learning inferences showed incursive *Ae. albopictus* had travelled to uninvaded regions from both adjacent and non-adjacent islands. Private alleles and a coancestry matrix indicated direct gene flow from PNG into nearby islands. Outlier analyses also detected four linked alleles introgressed from PNG, with the alleles surrounding 12 resistance-associated cytochrome P450 genes. By treating dispersal as both an intergenerational process and a set of discrete events, we describe a highly interconnected invasive system.

## Introduction

Population genetics has traditionally treated dispersal as an intergenerational process, but one that is nevertheless derived from the movement of individual organisms within each generation (Wright, 1943). Intergenerational dispersal describes how organisms distributed across geographical space are connected through time via a spatial pedigree (Bradburd & Ralph, 2019), and can be summarised at a population level by the mean distance between parent and offspring. Population genomic approaches applied to wild populations increasingly provide the power needed to detect dispersal across fine temporal scales down to single generations, which can reveal movement patterns at correspondingly fine spatial scales (Combs, Puckett, Richardson, Mims, & Munshi-South, 2018; Jasper, Schmidt, Ahmad, Sinkins, & Hoffmann, 2019; Trense et al., 2020). Spatial genomic studies conducted across a restricted temporal range may also help shed new light on adaptive processes such as the spread of advantageous alleles through wild populations (Endersby-Harshman et al., 2020; Fitzpatrick et al., 2010; Pélissié, Crossley, Cohen, & Schoville, 2018) and local adaptation in microgeographically structured environments (Yadav, Stow, & Dudaniec, 2020).

Individuals connected through recent generations of the spatial pedigree will be close kin, with full-sibs and half-sibs separated by a single generation and first cousins by two. Close kin dyads can be identified using genomics, and the spatial distribution of kin has been used in recent studies to detect dispersal over fine temporal scales (Combs et al., 2018; Escoda, Fernández-González, & Castresana, 2019; Escoda, González-Esteban, Gómez, & Castresana, 2017; Fountain et al., 2018; Jasper et al., 2019; Schmidt, Filipović, Hoffmann, & Rašić, 2018; Trense et al., 2020). Dispersal inferences from close kin treat dispersal as a set of discrete events reflecting specific acts of individual movement, making them particularly valuable for investigating dispersal through regions of genetic similarity, such as where populations have only recently become isolated (Escoda et al., 2019, 2017) or where a population has been sampled continuously across a range (Combs et al., 2018; Jasper et al., 2019; Schmidt et al., 2018; Trense et al., 2020). As kinship-based approaches assess recent movement, they are ideally applied to systems which have been subject to recent change, such as biological invasions or threatened species in disturbed habitats.

While the past 1-2 generations of dispersal represent very recent movement, population genomics has also been used to detect intragenerational dispersal, which has traditionally been investigated by directly observing the movement of individuals (Harrington et al., 2005; Schultz & Crone, 2001). Examples of this approach include tracing individual invasive species incursions (“incursives”) to their population of origin (M. Z. Chen et al., 2020; Schmidt, Chung, van Rooyen, et al., 2020; Schmidt et al., 2019). Although such studies require sampling across broad scales they are a useful way of detecting long-distance dispersal, which is difficult to investigate but can have important evolutionary consequences (Gillespie et al., 2012; Waters, Fraser, & Hewitt, 2013). Note that when incursive individuals are intercepted, there is no ‘dispersal’ in the intergenerational sense, but these movements are nevertheless important components of life-histories and food webs (Howard, 1960).

Recent human-mediated biological invasions are complex spatial processes that require careful investigation. In recent invasions, high regional coancestry throughout the invaded region will make the analysis of genetic structure alone insufficient for evaluating movement (Fitzpatrick, Fordyce, Niemiller, & Reynolds, 2012). Additionally, invasive species, defined here as any taxon which rapidly spreads (following Cristescu (2015)), frequently exhibit stratified dispersal, in which short-range active movement by the organism and long-range passive transportation by humans operate together across a range of spatial scales within a generation (Hengeveld, 1989; Sharov & Liebhold, 1998). Long-range dispersal can be traced within invaded regions or from distant origins (Schmidt, Chung, Honnen, Weeks, & Hoffmann, 2020; Sherpa et al., 2019), and can operate alongside short-range dispersal to spread adaptive alleles into and through invasive populations (Endersby-Harshman et al., 2020; Pélissié et al., 2018). Applying intragenerational and kinship-based dispersal methods to recent invasions may help detect recent movement across fine and broad spatial scales, with dispersal between invaded regions revealed through close kin and intragenerational incursions into uninvaded regions traced to their source.

Here we use spatial population genomics to investigate an invasion of *Aedes albopictus* (Asian tiger mosquito) in the Torres Strait Islands (TSI), Australia, sampled ∼14 years after colonisation. The Torres Strait lies between the northernmost point of the Australian mainland (Cape York) and Papua New Guinea (PNG) (Fig 1 inset). The region contains over 100 islands, of which 18 are inhabited communities (http://www.tsra.gov.au/the-torres-strait/community-profiles#TS%20Communities) and which we refer to here as ‘villages’ to distinguish from ecological communities. *Aedes albopictus* was first detected in the TSI in 2004 (Ritchie et al., 2006), and an insecticide-focussed elimination program began shortly thereafter. Logistical difficulties and recurrent reinvasion risks led to the abandonment of this program in 2008 in favour of a containment strategy, in which a *cordon sanitaire* was imposed to regulate the movement of people and goods between the invaded islands of the Torres Strait and the uninvaded ‘inner’ islands of Ngurapai (Horn) and Waiben (Thursday) (Muzari et al., 2017; van den Hurk et al., 2016). Ngurapai and Waiben serve as commercial hubs connecting the TSI and the Australian mainland, and the *cordon sanitaire* approach has likely helped prevent incursion of *Ae. albopictus* onto the mainland of Australia through the Torres Strait route though it continues to vector dengue outbreaks in the TSI (Muzari et al., 2017). *Aedes albopictus* incursions continue to be detected at ports on the Australian mainland and have been traced to locations in East and Southeast Asia that have strong trade links to Australia (Schmidt, Chung, van Rooyen, et al., 2020). Previous investigations of TSI *Ae. albopictus* using mtDNA and microsatellites indicated high coancestry with Indonesian *Ae. albopictus* (Beebe et al., 2013; Maynard et al., 2017). Additional observations of spatially and temporally variable genetic structure within the TSI could reflect high regional gene flow following local founder events (Maynard et al., 2017).

**Figure 1:**
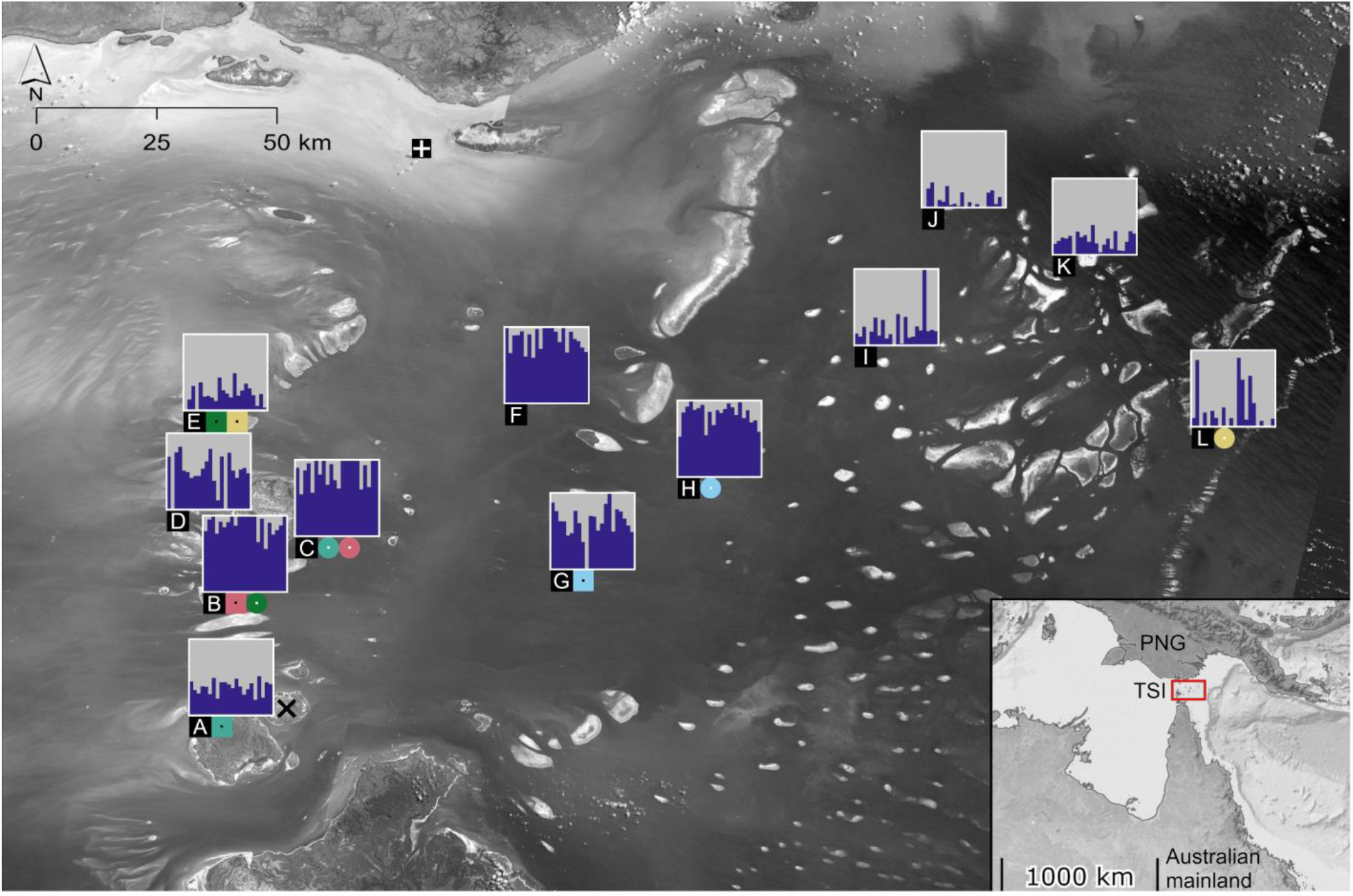
Locations and genetic structure of the TSI villages. A:Keriri, B:St Pauls (Moa Island) C:Kubin (Moa Island), D:Badu, E:Mabuiag, F:Iama, G:Warraber, H:Poruma, I:Masig, J:Ugar, K:Erub, L:Mer. Plots are of sparse non-negative matrix factorisation (sNMF) on the Region22 dataset (see Datasets and filtering), and are centred over each village. Grey and blue sections of each vertical bar represent ancestral lineages of individuals, assuming K = 2. White-dotted circles and black-dotted squares of the same colour indicate close kin dyads found across villages, with squares denoting the origin (see Table 2). The white crossed square indicates Dauan. The “**×**” indicates Ngurapai, where the four incursive mosquitoes were detected. Map inset shows the Australian mainland, Papua New Guinea (PNG) and the Torres Strait Islands (TSI). See Figures S1-S13 for village maps and sampling point locations, and Figures S15-S17 for sNMF results for 2 ≤ K ≤ 4.

While Indonesia is the likely source of the initial invasion into the TSI, over 5,000 boat journeys are made each year between TSI and PNG (Mcfarlane, 1998), indicating the potential for gene flow across this border. When gene flow takes place after an initial invasion it can operate as a ‘genetic invasion’ if it interferes with control strategies, such as through introgression of alleles conferring insecticide resistance (Endersby-Harshman et al., 2020; Riveron et al., 2013; Schmidt et al., 2019). Although the TSI was likely invaded from Indonesia via a ‘stepping-stone’ in the Southern Fly region of PNG (Beebe et al., 2013), the absence of *Ae. albopictus* in Southern Fly in the late 1990s (Johansen et al., 2000) and its present genetic similarity to the TSI (Maynard et al., 2017) indicates that the invasion of the TSI and Southern Fly regions happened contemporaneously from Indonesia. PNG *Ae. albopictus* from outside the Southern Fly region are genetically distinct from Indonesian *Ae. albopictus* (Maynard et al., 2017) and gene flow from this PNG background should be readily detectable.

This investigation covers a range of spatial scales from 10s of metres to 1000s of kilometres, and temporal scales from a single generation to ∼100 generations in the past. Within the TSI, where coancestry is high, we focus on dispersal in the immediate past, covering both short-range active flight within islands and long-range passive transportation between islands, including incursions past the *cordon sanitaire*. At broader spatial scales, we use a panel of differentiated genotypes from other locations including nearby Papua New Guinea (PNG) to investigate earlier patterns of gene flow into the TSI. Our specific aims are: (i) to investigate the spatial structure of passive dispersal, and whether it occurs only between nearby locations or distant locations as well; (ii) to ascertain whether new genetic structure among islands has developed since previous investigation (Maynard et al., 2017); and (iii) to determine whether gene flow has occurred from PNG into the TSI, including the spread of putatively adaptive alleles. Through this, we show how stratified dispersal operates in this system and how long-range dispersal can assist the rapid invasion of new regions and the spread of advantageous alleles through established populations. The rapid spread of *Ae. albopictus* and its capacity for long-distance invasion make studies of its dispersal globally relevant (Goubert et al., 2017; Schmidt, Chung, Honnen, et al., 2020; Sherpa et al., 2019).

## Materials and Methods

### Sample acquisition and genotyping

We analysed *Ae. albopictus* single nucleotide polymorphism (SNP) data from multiple sources. These included new mosquito samples collected from the Torres Strait Islands (TSI) and Papua New Guinea (PNG) as well as sequence data collected previously from across the Indo-Pacific region. We use the alphabetical key (A-L) in Fig 1 in all references to TSI villages.

Sampling in the invaded regions of the TSI took place at two time points. The first involved cross-sectional collections from 13 villages on 12 islands sampled between 2018-04-24 and 2018-05-04 (Fig 1). On Moa Island, two villages were sampled (B:St Pauls and C:Kubin). By restricting the temporal range of sampling, it is possible to undertake investigations within and between villages that require data at a fine temporal scale (i.e. within 1–2 generations), such as kinship analysis. Within each village, researchers selected and georeferenced sampling points where they collected adult *Ae. albopictus* with sweep-nets. Sampling points were separated by at least 100 m where possible (maximum 3,297 m), and each village had 15–20 sampling points except Dauan which had 7 (Fig 1: red crossed square). *Aedes albopictus* were preserved in 100% ethanol prior to DNA extraction. A second TSI sampling event took place on I:Masig between 2019-03-28 and 2019-04-08 at the same sampling points, serving as a temporal replicate. Village maps and sampling point locations are detailed in Figures S1-S13.

In February 2019, we detected *Ae. albopictus* incursions on the island of Ngurapai, south of the *cordon sanitaire* (see Fig 2). Four incursive *Ae. albopictus* adults were detected between 2019-02-02 and 2019-02-09 during routine wet season surveillance (Figure S14) and the samples were preserved for DNA extraction. Similar surveys, conducted with sweep nets in December 2018, March 2019 and May 2019 did not detect any *Ae. albopictus* on the island.

**Figure 2:**
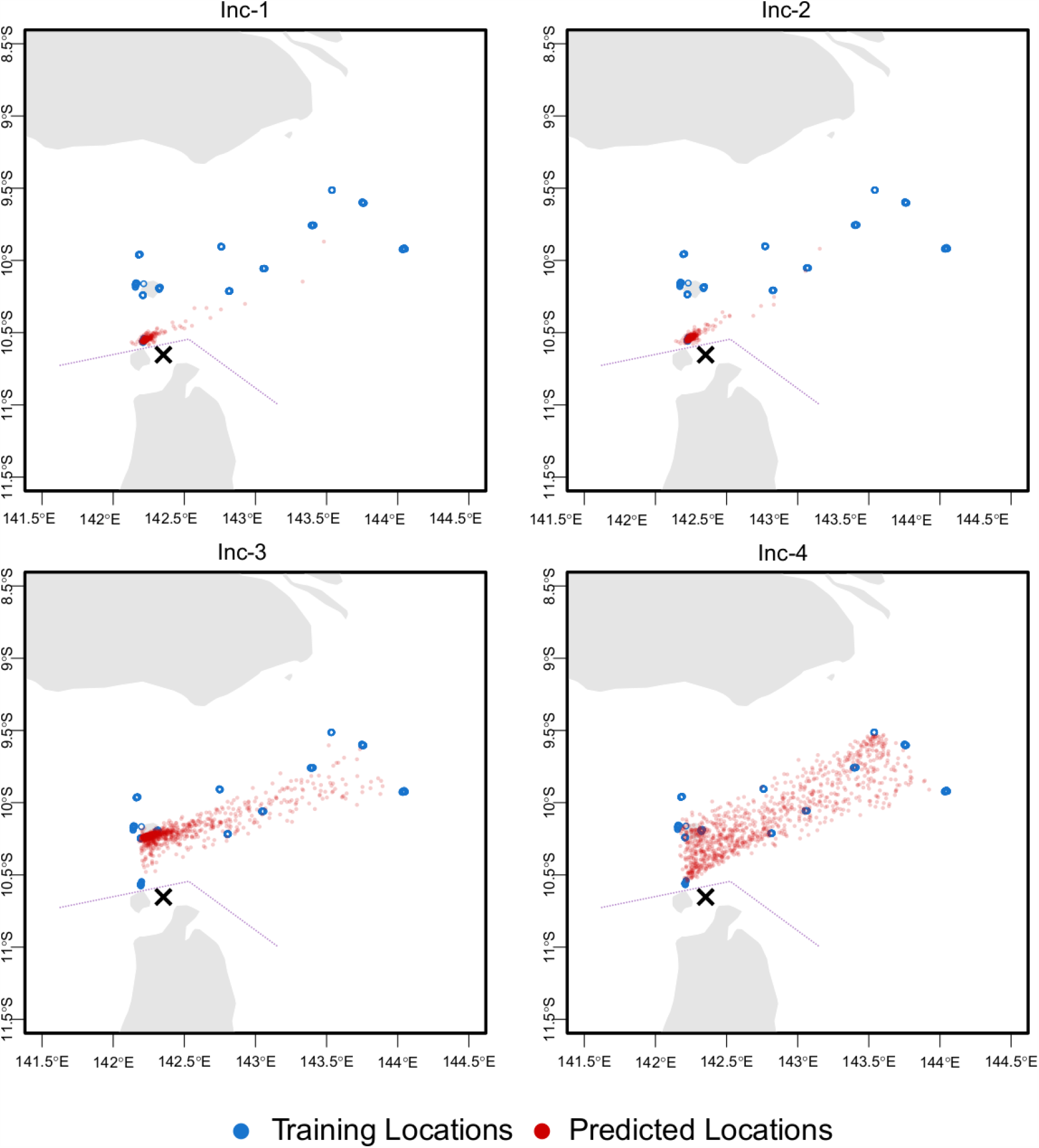
Predicted locations of the four incursives from 1000 bootstrapping runs of Locator. Each prediction is depicted with a red circle. The “**×**” indicates Ngurapai, where the incursives were detected. The dotted line describes the position of the *cordon sanitaire*. See main text for point estimates of locations.

A final set of samples was collected from PNG between 2018-08-02 and 2018-09-13 using ovitraps deployed at several urban locations around the cities of Port Moresby (south coast) and Madang (north coast). These were aggregated to produce one Port Moresby sample and one Madang sample, with a maximum of two mosquitoes taken from each individual ovitrap to reduce the likelihood of sampling close kin.

Samples were genotyped using a pipeline described previously (Schmidt, Chung, van Rooyen, et al., 2020) and described in full in Text S1. Briefly, we used the double digest restriction site-associated (ddRAD) sequencing protocol developed for *Ae. aegypti* (Rašić, Filipović, Weeks, & Hoffmann, 2014) and validated in *Ae. albopictus* (Schmidt et al., 2017). Sequence reads were processed in Stacks v2.41 (Catchen, Hohenlohe, Bassham, Amores, & Cresko, 2013), with Bowtie2 v2.3.5.1 (Langmead & Salzberg, 2012) used to align reads to the AalbF2 genome assembly (Palatini et al., 2020).

Sequence data were generated for 371 *Ae. albopictus*: 301 from the cross-sectional collections on the invaded islands, 30 from the temporal replicate on I:Masig, 4 from the incursions past the *cordon sanitaire*, and 18 from each of Madang and Port Moresby in PNG. We also included sequence data from previous work (Schmidt, Chung, Honnen, et al., 2020; Schmidt, Chung, van Rooyen, et al., 2020). This included sequences from Indonesia (Bali, Bandung and Jakarta) and Timor-Leste, the proposed approximate origin of the initial TSI invasion (Beebe et al., 2013; Maynard et al., 2017). We also added sequences from locations linked to Australian *Ae. albopictus* incursions: China (Guangzhou), Japan, Singapore and Taiwan (Schmidt, Chung, van Rooyen, et al., 2020). Finally, we included two Pacific Island locations from East of PNG: Fiji and Vanuatu. These data were all reprocessed through the same bioinformatics pipeline.

### Datasets and filtering

Genotypes were filtered with the Stacks v2.41 program Populations (Catchen et al., 2013), VCFtools v0.1.16 (Danecek et al., 2011), and Beagle v4.1 (Browning & Browning, 2016). Populations produced VCF files containing SNPs called in ≥ 0.75 of the mosquitoes from each village, location or timepoint; VCFtools retained biallelic SNPs with minor allele count ≥ 3 (Linck & Battey, 2019), depth of coverage ≥ 5, and genotyped in ≥ 80% of total mosquitoes; and Beagle imputed and phased genotypes in 50,000 bp sliding windows with 3,000 bp overlap.

We first used Populations and VCFtools to detect close kin dyads (1^st^ and 2^nd^ order) within each village. We used the *relatedness2* command in VCFtools to detect first-order (parental or full-sib) and second-order (grandparental or half-sib) kin dyads, analysing each village separately. This command uses the KING (Manichaikul et al., 2010) method of generating kinship coefficients for pairwise relationship inference; accordingly, we used kinship coefficient cut-offs of > 0.177 for first-order kin and > 0.088 for second order kin. Close kin dyads identified at this step were used to help estimate kinship categories of dyads across villages (see ‘Close kin across villages’ below).

For analysis within the TSI, we were cautious to avoid filtering bias caused by close kin and uneven sample sizes (Puechmaille, 2016). Accordingly, we produced datasets of n = 22 for each village, with 22 the maximum number of genotypes that ensured equal sample sizes after removing close kin within and across villages (see Results). Kin were removed in order of missing data, other genotypes were removed to maximise the geographical distribution of genotypes, then by missing data. Datasets were imputed and phased with Beagle and hereafter called the “Village22” datasets, each containing between 7,877–9,873 SNPs.

We also produced a dataset for the TSI region called “Region22” that contained all individuals from the Village22 datasets, containing 26,085 SNPs. Tracing of incursions and gene flow from PNG used this dataset where specified, with the incursive or PNG genotypes included. For analyses using the additional Indo-Pacific genotypes, we included all non-incursive individuals from the TSI including Dauan, with the temporal replicate from I:Masig excluded and with the dataset left unimputed where indicated. This dataset contained 495 mosquitoes genotyped at 26,970 SNPs.

### Population processes within the Torres Strait

#### TSI genetic structure

We investigated genetic structure in the TSI using sparse non-negative matrix factorisation (sNMF) in the R package “LEA” (Frichot & François, 2015), run on the Region22 dataset. This analysis estimates individual ancestry coefficients, assuming that individual genotypes are produced from the admixture of K ancestral lineages. Previous research using microsatellites to analyse samples from 2007–2015 found two main genetic lineages in the TSI (Maynard et al., 2017). Accordingly, we ran 50 iterations of the sNMF with 2 ≤ K ≤ 4 but used K = 2 for visualisations to provide a direct comparison with this work.

We used Populations to calculate heterozygosity (H_O_) and nucleotide diversity (π) at variant sites and pairwise F_ST_ between all village samples. We used VCFtools to calculate Tajima’s *D* in 10 Mbp windows, including the temporal replicate from I:Masig to reveal any changes over the 12 month period. We used the R package “hierfstat” (Goudet, 2005) to estimate global F_ST_ and the population differentiation statistic D_est_ (Jost, 2008) among the 2018 samples.

#### Dispersal within villages

Within TSI villages (< 3.5 km), dispersal is mostly by active flight through continuous habitat and can be summarised by the parameter σ, which makes up the dispersal variance component of neighbourhood size (N_W_: Wright, 1946). N_W_ can be thought of as the effective number of *Ae. albopictus* that make up a mosquito’s breeding ‘neighbourhood’ when isolation by distance is operating. The equation, N_W_ = 4π. σ^2^. d, provides a link between spatial patterns of genetic differentiation and local demographic processes, connecting intergenerational dispersal (σ^2^) to the effective density of breeding adults within the dispersal area (d).

We used the Village22 datasets to investigate isolation by distance within each village. Mantel tests (function mantel.randtest) ran in the R package “ade4” (Dray & Dufour, 2007) analysed matrices of individual linear genetic distance and the natural logarithm of Haversine geographic distance, using 9,999 permutations and Bonferroni correction to assess statistical significance. Rousset’s *a* (Rousset, 2000) provided genetic distance, calculated in SPAGeDI (Hardy & Vekemans, 2002).

We estimated neighbourhood size (N_W_: Wright, 1946) within all villages using the inverse of the regression slope of pairwise genetic distance (Rousset’s *a*) against the natural logarithm of geographical distance (Rousset, 2000). As the linearity of this relationship may break down at distances within the dispersal estimate σ, we omitted pairs < 50 m apart. The temporal replicate from I:Masig was also omitted. The pairwise data used for Mantel tests were concatenated to run a single linear regression on the 5325 within-village dyads.

#### Close kin across villages

Having identified close kin dyads within villages, we applied a conservative process for detecting kin across villages, which represent passive transportation by humans. First, we generated kinship coefficients using PC-Relate (Conomos, Reiner, Weir, & Thornton, 2016), which controls for genetic structure by conditioning the data with principal components (PCs). We ran PC-Relate on the dataset containing all non-incursive individuals, first pruning SNPs by linkage disequilibrium (R package “SNPRelate” (Zheng et al., 2012); using the snpgdsLDpruning function with arguments, method = “composite” and ld.threshold = 0.2).

We generated kinship coefficients for all dyads within and across villages following three different conditioning treatments: 3 PCs, 5 PCs, and 10 PCs. As a lower bound for determining which dyads across villages could be considered to have a particular category of relatedness, we used the lowest kinship coefficient observed among all close kin dyads within villages (see Datasets and filtering).

For each dyad found across two villages, one individual (Sib-1) or its recent relative will have dispersed to a new environment (Village-1) while the other (Sib-2) has stayed at the origin (Village-2). Thus, Sib-1 and Sib-2 should both have higher coancestry with individuals from Village-2 than those from Village-1, which should be reflected in higher average kinship with Village-2. We estimated the direction of dispersal by comparing kinship coefficients of Sib-1 and Sib-2 with each village, which allowed us to determine the origin (Village-2) and the destination (Village-1). These differences were evaluated with t-tests.

### Incursions past the *cordon sanitaire*

To estimate source locations of the four incursive *Ae. albopictus*, we used two complementary methods in the programs assignPOP (K.-Y. Chen et al., 2018) and Locator (Battey, Ralph, & Kern, 2020). assignPOP treats each village as a population, and generates assignment probabilities to each hypothetical population using a Monte Carlo assignment test with a support vector machine predictive model. Locator makes no population-based assumptions, but uses geolocations of individual genotypes to infer the spatial origin of each incursive using deep learning. These analyses used the Region22 dataset, omitting the temporal replicate from I:Masig.

Following Schmidt et al. (2019), we used the assignPOP function assign.X to generate posterior probabilities of assignment from which we calculated ‘relative probabilities’ of assignment, defined as the highest posterior probability divided by the second highest. Relative probability > 3 has previously been used as a cut-off for *Ae. albopictus* assignment at broad scales (Schmidt, Chung, van Rooyen, et al., 2020).

To run Locator we first pruned SNPs by linkage disequilibrium using the snpgdsLDpruning function in the R package “SNPRelate” (arguments, method = composite and ld.threshold = 0.2). This dataset provided a point estimate of each incursive’s original location, and 1000 bootstrap subsamples were run to provide confidence around these estimates. While Locator has options for analysing SNPs in windows, our ddRADseq data were too sparse and the reference assembly insufficient for this.

Before running final assignments, we first confirmed that each of the four incursives had an origin in the TSI through an initial run of assignPOP using the dataset containing all non-incursive Indo-Pacific genotypes. These results showed strong support for each incursive mosquito having an origin in the TSI, with aggregated posterior assignment probabilities of 0.89–0.97 to the TSI and 0.11–0.03 to non-TSI locations (Table S1).

### International gene flow into the Torres Strait

#### Genome-wide gene flow

We used fineRADstructure (Malinsky, Trucchi, Lawson, & Falush, 2018) to investigate possible coancestry between TSI *Ae. albopictus* and those from locations other than Indonesia or Timor-Leste. We ran fineRADstructure with default settings on the dataset containing all non-incursive Indo-Pacific genotypes, with the temporal replicate of I:Masig removed.

This analysis also helped clarify genetic relationships among genotypes in the broader region. To assist with the visualisation of these relationships, we ran a PCA-UMAP analysis (Diaz-Papkovich, Anderson-Trocmé, Ben-Eghan, & Gravel, 2019) on the same dataset. PCA-UMAP used the first 4 PCs, projected in two dimensions via UMAP using 50 neighbours and a 0.5 minimum distance.

Using the Region22 dataset, we calculated the number of private alleles in each TSI village with Populations, excluding the temporal replicate from I:Masig. This was then recalculated with the two PNG samples (Port Moresby and Madang) included in filtering. By comparing the number of private alleles in each village with and without the PNG samples, we assessed gene flow between each village and PNG. Specifically, if fewer private alleles were recorded for a village after the PNG samples were included in filtering, the “missing” alleles will be identical by state and likely identical by descent in both PNG and that village. As these alleles are restricted to specific villages, they are evidence of gene flow from PNG into each village directly.

#### Adaptive introgression

While ddRAD data are typically too sparse to detect alleles under selection, they can be suitable for detecting selective sweeps as shown by previous work on *Ae. aegypti* (Endersby-Harshman et al., 2020). We looked for signs that advantageous alleles had spread into the TSI using a three-step process: identifying genomic regions of interest, where multiple SNPs had irregular structure consistent with linked selection; identifying genes within these genomic regions of potential adaptive importance; and identifying the geographical origin of alleles at these SNPs.

We used the R packages *pcadapt* (Luu, Bazin, & Blum, 2017) and LEA (Frichot & François, 2015: function sNMF) to detect SNPs with irregular structure. Analyses were run on the Region22 dataset. We ran *pcadapt* with a minimum allele frequency of 0.025 (≥ 15 allele copies present), and obtained P-values from Mahalanobis distances. Genomic inflation factors for all runs were < 1.5. We ran sNMF using default parameters and 50 repetitions, and selected outliers by F_ST_. For each analysis, P-values were transformed using a Bonferroni correction, and we used a q-value cut-off of 0.001. Only SNPs detected as outliers by both *pcadapt* and sNMF were considered. As genotype imputation methods can introduce bias when imputing rare alleles (Shi et al., 2018), we repeated the above methods without imputation in Beagle.

Within TSI *Ae. albopictus*, local demographic changes such as founder effects may have eliminated genetic diversity in particular regions of the genome, which a genome scan may incorrectly read as the result of a selective sweep (Hoban et al., 2016). For this reason, we restricted our investigation to outlier regions that satisfied three conditions: (i) the region contained multiple outlier SNPs spread across multiple RADtags within 1 Mbp; (ii) the region contained one or more genes with products of plausible adaptive importance, such as insecticide resistance; and (iii) the alleles at the specific outlier SNPs within the region were not found in Indonesia (putative source of the TSI invasion) but were found in other populations outside of the TSI. For (ii) we used Integrative Genomics Viewer (Robinson et al., 2011) to visually explore the *Ae. albopictus* reference assembly. For (iii), we used the dataset containing all non-incursive Indo-Pacific genotypes (N = 525), without imputation.

## Results

### Population processes within the Torres Strait

#### TSI genetic structure

Analysis of genetic structure with sNMF provided a picture of genetic patterns across the TSI for K = 2 (Figure 1). Despite the higher resolution of SNP markers and the additional ∼3-11 years elapsed since sampling, there was no clearer spatial genetic structure within and between villages at K = 2 than in the previous microsatellite study (c.f. Maynard et al., 2017). Both ancestral lineages were detected in every village, though 48 individuals were assessed as not admixed (Figure 1). Genetic admixture among individuals was highly variable in some villages, particularly in D:Badu, G:Warraber, I:Masig and L:Mer. Despite this, some geographical patterns were evident from the K = 2 analysis, particularly the genetic similarity of the central islands (F:Iama, G:Warraber, H:Poruma) and those of Moa Island (B:St Pauls and C:Kubin). These patterns are increasingly pronounced at K = 3 (Figure S16), though the high admixture within villages (Figures 1,S15,S16,S17) and low pairwise F_ST_ among villages (Table S2) indicate that genetic structure is weak within the TSI. Genetic similarity among the central islands has previously been noted (Maynard et al., 2017), and points to temporal consistency in genetic structure.

Mean global F_ST_ among villages in 2018 was 0.039 (SE = 0.000) while differentiation (D_est_) was 0.006 (SE = 0.000). Pairwise F_ST_ estimates between villages (Table S2) indicate that I:Masig had higher average pairwise F_ST_ in 2019 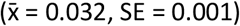 than in 2018 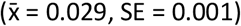, an average increase in F_ST_ of 8.9% over the 12 months, perhaps reflecting the development of additional genetic structure across time. Tajima’s D can be difficult to interpret, but the consistently negative mean values across all villages accord with past population expansions following colonisation (Gattepaille, Jakobsson, & Blum, 2013).

#### Dispersal within villages

Table 1 lists mean pairwise genetic distance (Rousset’s *a*) for dyads sampled at < 50 m and > 500 m separation within each village. Mosquitoes sampled at > 500 m separation did not have consistently higher genetic distances than those sampled at < 50 m separation. Likewise, 6 of the 13 Mantel tests showed a negative relationship between genetic and geographic distance within villages (Figures S18-S30). There was no statistically significant isolation by distance within any of the villages (all Bonferroni-corrected P-values > 0.05).

**Table 1:** TSI village details and population genetic parameter estimates including heterozygosity (H_O_) and nucleotide diversity (π) at variant sites, Tajima’s D, Rousset’s *a* for dyads < 50 m and > 500 m apart, and number of private alleles with and without inclusion of PNG samples.

| Village key (Fig 1) | Village | Island area (km<sup>2</sup>) | Human pop size (2016) | $H_o$ (±SE) | $\pi$ (±SE) | Tajima's D (±SE) | Mean (±SD) Rousset's $a$: <50m | Mean (±SD) Rousset's $a$: >500m | Private alleles (no PNG) | Private alleles (PNG) |
| --- | --- | --- | --- | --- | --- | --- | --- | --- | --- | --- |
| A | Keriri | 14.5 | 268 | 0.063 (±0.001) | 0.074 (±0.001) | -1.164 (±0.026) | 0.25 (±0.07) | 0.20 (±0.10) | 118 | 98 |
| B | St Pauls (Moa) | 152.4 | 248 | 0.063 (±0.001) | 0.074 (±0.001) | -1.010 (±0.030) | 0.18 (±0.08) | 0.20 (±0.10) | 123 | 91 |
| C | Kubin (Moa) | 152.4 | 198 | 0.064 (±0.001) | 0.075 (±0.001) | -0.809 (±0.032) | 0.16 (±0.09) | 0.16 (±0.08) | 97 | 80 |
| D | Badu | 101.9 | 813 | 0.065 (±0.001) | 0.078 (±0.001) | -1.164 (±0.028) | 0.18 (±0.07) | 0.21 (±0.09) | 56 | 42 |
| E | Mabuiag | 6.3 | 210 | 0.065 (±0.001) | 0.076 (±0.001) | -0.986 (±0.028) | 0.15 (±0.11) | 0.19 (±0.06) | 146 | 115 |
| F | Iama | 1.5 | 319 | 0.063 (±0.001) | 0.074 (±0.001) | -0.691 (±0.034) | 0.21 (±0.08) | 0.12 (±0.07) | 120 | 85 |
| G | Warraber | 0.6 | 245 | 0.066 (±0.001) | 0.078 (±0.001) | -0.980 (±0.027) | 0.20 (±0.06) | 0.23 (±0.05) | 77 | 66 |
| H | Poruma | 0.4 | 167 | 0.065 (±0.001) | 0.076 (±0.001) | -0.972 (±0.028) | 0.17 (±0.08) | 0.19 (±0.07) | 105 | 83 |
| I | Masig (2018) | 1.5 | 270 | 0.069 (±0.001) | 0.080 (±0.001) | -1.054 (±0.027) | 0.11 (±0.00) | 0.19 (±0.05) | 128 | 90 |
| I | Masig (2019) | | | 0.073 (±0.001) | 0.082 (±0.001) | -0.908 (±0.029) | 0.14 (±0.02) | 0.14 (±0.04) | | |
| J | Ugar | 0.4 | 85 | 0.063 (±0.001) | 0.074 (±0.001) | -1.106 (±0.029) | 0.22 (±0.06) | 0.19 (±0.05) | 130 | 85 |
| K | Erub | 6.9 | 328 | 0.063 (±0.001) | 0.076 (±0.001) | -1.209 (±0.027) | 0.19 (±0.09) | 0.24 (±0.08) | 182 | 123 |
| L | Mer | 4.3 | 450 | 0.067 (±0.001) | 0.081 (±0.001) | -1.240 (±0.026) | 0.22 (±0.10) | 0.25 (±0.09) | 148 | 77 |

When within-village dyads were concatenated to estimate neighbourhood size (N_W_), isolation by distance was revealed by the positive relationship between genetic distance (Rousset’s *a*) and the natural logarithm of geographical distance (geodist): *a* = 0.003025 × geodist + 0.1878, with coefficient SE = 0.00129. N_W_ within villages was estimated at 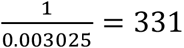 mosquitoes (95% C.I. 232–577). Despite high variability among villages, these results indicate that TSI *Ae. albopictus* have some spatial structure at scales of 100s to 1000s of metres.

#### Close kin within villages

Three of the 2772 dyads within villages had putative first-order relatedness, and none had second-order relatedness (Table 2). Two of the three dyads were collected at the same sampling point, the other at points 170 m apart. This low incidence of close kin compared with other studies (e.g. Jasper et al., 2019) was likely due to the 100 m separation between most sampling points and suggests mostly local movement within villages.

**Table 2:** Putative close kin dyads within and across villages. Within villages, the dyad at E:Mabuiag had the lowest PC-Relate kinship coefficient of the three close kin dyads, so this was used as the cut-off for designating close kin across villages. Mean kinship describes the mean of kinship coefficients between Sib-1 and Sib-2 and all other individuals from that village, using kinship coefficients from the 3 PC treatment. Higher mean kinship in Village-2 shows this is the origin.

| Sib-1 | Sib-2 | Origin | Distance (km) | Kinship (KING) | Kinship (PC-Relate 3 PCs) | Kinship (PC-Relate 10 PCs) | Mean kinship Village-1 (±SD) | Mean kinship Village-2 (±SD) |
| --- | --- | --- | --- | --- | --- | --- | --- | --- |
| **Within villages** |  |  |  |  |  |  |  |  |
| K:Erub | K:Erub |  | 0.17 | 0.43 | 0.59 | 0.47 |  |  |
| B:St Pauls | B:St Pauls |  | 0 | 0.42 | 0.51 | 0.48 |  |  |
| E:Mabuiag | E:Mabuiag |  | 0 | 0.34 | 0.32 | 0.28 |  |  |
| **Between villages** |  |  |  |  |  |  |  |  |
| C:Kubin | A:Keriri | A:Keriri | 35.26 |  | 0.52 | 0.35 | -0.015 (±0.017) | 0.042 (±0.038) |
| H:Poruma | G:Warraber | G:Warraber | 31.51 |  | 0.51 | 0.41 | -0.017 (±0.017) | 0.032 (±0.033) |
| C:Kubin | B:St Pauls | B:St Pauls | 13.50 |  | 0.49 | 0.46 | -0.036 (±0.018) | 0.001 (±0.021) |
| L:Mer | E:Mabuiag | E:Mabuiag | 202.63 |  | 0.44 | 0.42 | -0.013 (±0.018) | 0.055 (±0.052) |
| B:St Pauls | E:Mabuiag | E:Mabuiag | 30.59 |  | 0.44 | 0.44 | -0.002 (±0.016) | 0.060 (±0.040) |

#### Close kin across villages

Within villages, the dyad in E:Mabuiag had the lowest PC-Relate kinship coefficient of the three close kin dyads (0.32 with 3 PCs; Table 2), so this was used as the cut-off for designating close kin across villages. Five of the 31,944 dyads across villages had higher scores than this, indicating putative first-order relatedness. The three highest kinship coefficients for ‘unrelated’ dyads were 0.27, 0.23, and 0.16. When 5 or 10 PCs were included, the rank order of kinship coefficients fluctuated but all remained consistently above the within-village cut-off and above the coefficients of unrelated dyads.

Four of the putative close kin dyads were across nearby villages while the fifth was between E:Mabuiag and L:Mer, ∼200 km apart (Figure 1, Table 2). For each dyad the direction of dispersal was apparent, demonstrated by both individuals in the dyad having higher mean kinship coefficients with Village-2 (the origin) than Village-1 (the destination). Results of t-tests assessing dispersal direction were all strongly statistically significant (all t > 8.6, all P < 0.001).

### Incursions past the *cordon sanitaire*

The two methods of assignment, assignPOP and Locator, provided concordant results for two of the incursives (Fig 2). Inc-1 and Inc-2 had both clearly dispersed from A:Keriri, less than 10 km from where they were detected on Ngurapai (Fig 2: **×**). In assignPOP this was demonstrated by the high posterior (0.90 and 0.77) and relative (61.3 and 16.8) probabilities of assignment to A:Keriri. For Locator each incursive was placed among the A:Keriri genotypes for the clear majority of the 1000 bootstrap subsamples (Fig 2). Point estimates from Locator placed Inc-1 (10.5562° S, 142.2197° E) and Inc-2 (10.5585° S, 142.2173° E) amidst the A:Keriri samples.

Inc-3 and Inc-4 were assigned differently by assignPOP and Locator. This was not likely due to missing data (3.8% and 8.3%), as successful assignments with assignPOP have been recorded with > 40% missing data (Schmidt et al., 2019). assignPOP selected distant L:Mer (Inc-3) and I:Masig (Inc-4) as the most likely sources, though posterior (0.35 and 0.16) and relative (2.8 and 1.1) probabilities were low. Locator point estimates placed Inc-3 (10.2086° S, 142.2910° E) on Moa Island between the B:St Pauls and C:Kubin genotypes. Inc-4 (9.9216° S, 143.1598° E) was placed in the Torres Strait near I:Masig. Bootstrap subsampling revealed higher uncertainty for Inc-3 than for Inc-1 and Inc-2 but nevertheless indicated Moa Island was the likely source. No clear assignment was forthcoming for Inc-4.

### International gene flow into the Torres Strait

#### Genome-wide gene flow

The *fineRADstructure* plot (Fig 3) shows coancestry between genotype pairs from TSI and other selected locations. See Figure S31 for the figure at full size, and Figure S32 for a PCA-UMAP ordination of the same genotypes using the same colour symbology. TSI genotypes were only weakly clustered by location, as seen by the poor ‘sorting’ of genotypes in the colour panel. Only one village (J:Ugar) had every genotype resolved into a single lineage. However, TSI genotypes formed a lineage with Indonesia and Timor-Leste (Fig 3: solid black square, top right) distinct from all other genotypes (solid black square, bottom left).

**Figure 3:**
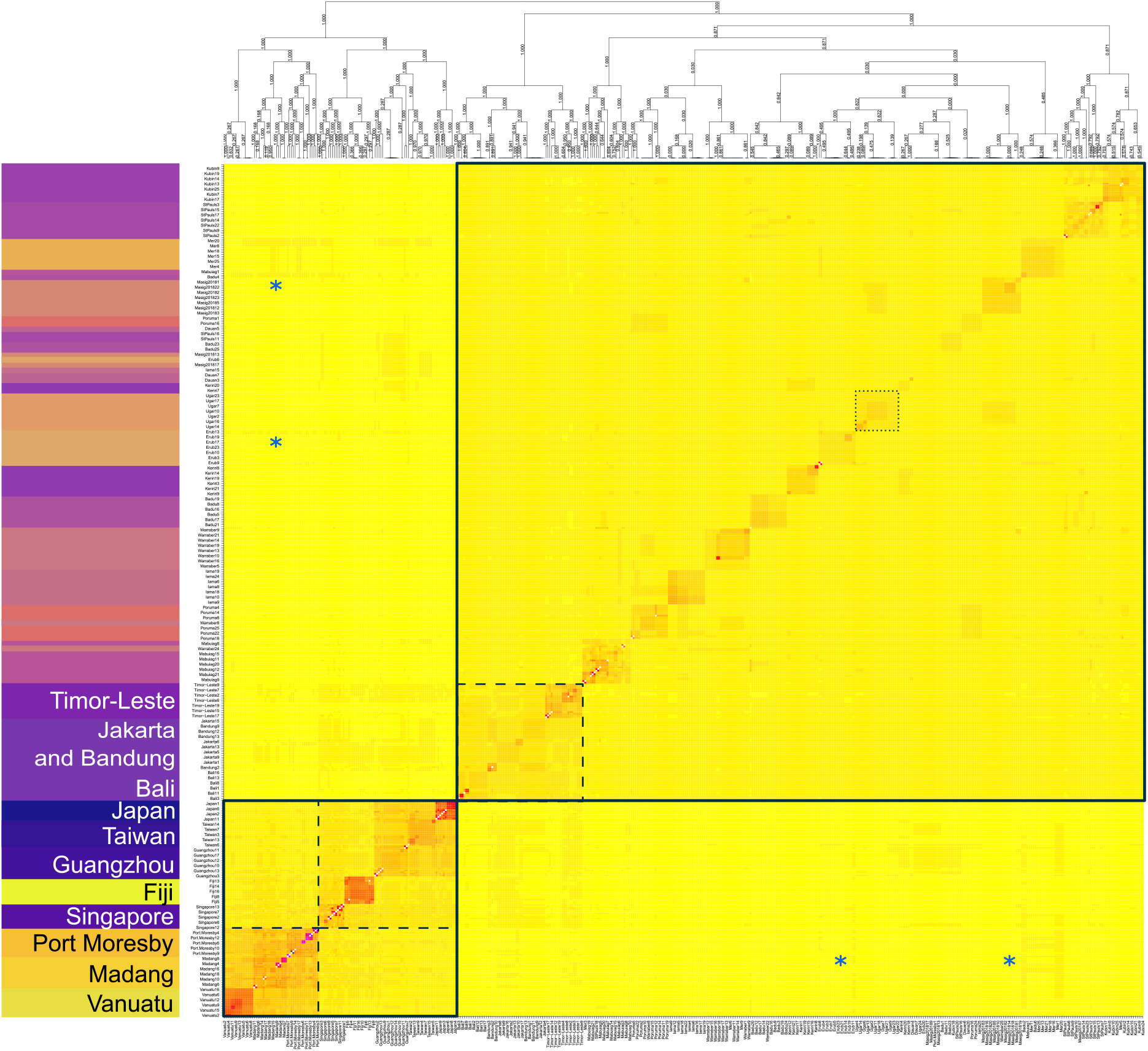
FineRADstructure coancestry map and tree. The left hand side panel indicates genotype sampling locations coloured by geographical proximity and with all non-TSI genotypes labelled. The central panel shows coancestry between genotypes, with light yellow indicating low coancestry, and darker yellows, oranges and reds indicating progressively higher coancestry. Solid black squares indicate the two major clades of TSI/Indonesia/Timor-Leste (top right) and all other genotypes (bottom left). Dashed squares indicate subgroups of PNG/Vanuatu, East Asia/Southeast Asia/Fiji, and Indonesia/Timor-Leste. The dotted square indicates J:Ugar, the only TSI village resolved into a single lineage. Blue asterisks indicate higher coancestry between PNG and TSI genotypes, particularly those from L:Mer. See Figure S31 for full-size figure.

Within these broad groups were subgroups (Fig 3: dashed squares) placing East and Southeast Asian genotypes together with Fiji and Indonesian genotypes with Timor-Leste, as expected from previous analysis (Schmidt, Chung, Honnen, et al., 2020; Schmidt, Chung, van Rooyen, et al., 2020). PNG genotypes grouped with Vanuatu, suggesting recent coancestry. While the PNG and Vanuatu lineage had slightly greater coancestry with Southeast Asia than East Asia, this lineage was clearly differentiated from all others. PCA-UMAP confirmed the three major groups of TSI/Indonesia/Timor-Leste, East Asia/Southeast Asia/Fiji, and PNG/Vanuatu (Figure S32).

Gene flow from PNG into the TSI was suggested by the higher coancestry between PNG and some TSI genotypes, particularly those from L:Mer (Fig 3: blue asterisks). PNG also had higher coancestry with Jakarta and Bandung in Indonesia but not with Bali. While there was some evidence of coancestry between East Asian and TSI genotypes, this was also observed in East Asian and Indonesian genotypes, particularly those from Jakarta and Bandung. It was thus unclear whether recent gene flow had occurred from East Asia into the TSI or whether past gene flow from East Asia into Indonesia had introduced alleles that were then brought to the TSI during invasion.

Table 1 lists the number of private alleles in each village with and without the PNG samples included in filtering. When PNG was included, the greatest ‘loss’ of private alleles was observed in the eastern islands of I:Masig (30% loss), J:Ugar (35% loss), K:Erub (32% loss), and L:Mer (48% loss) and northern island of F:Iama (29% loss). These ‘lost’ alleles are identical by state and likely also identical by descent to those in PNG, indicating gene flow between each island and PNG. Notably, the fact that these alleles were not found on other islands suggests that this gene flow was also specific to each island, and thus is not due to one single incursion into the TSI followed by local spread.

#### Adaptive introgression

Outlier analysis revealed four genomic regions of interest, each containing irregularly structured SNPs spread across multiple RADtags within 1 Mbp, and detected as outliers by both *pcadapt* and sNMF. For pcadapt, a scree plot of the proportion of explained variance did not indicate an optimum K (Figure S33). However, visual inspection of scatterplots of the first eight PCs indicated that genetic structure among villages was much less apparent after the first four PCs (Figures S34-37). Accordingly, we restricted our analyses to 2 ≤ K ≤ 4 for both programs.

The four genomic regions and all genes of known function contained therein are listed in Tables S3-S6. One of these regions, consisting of four SNPs across two RADtags ∼330 kbp apart on scaffold NW_021838465.1, contained 12 genes of interest (Table S3). These included three cytochrome P450 9e2-like genes and nine probable cytochrome P450 9f2 genes. Cytochrome P450 9e2 genes have been linked to insecticide resistance and demonstrated strong upregulation in resistant *Ae. aegypti* mosquitoes (Kim Lien et al., 2019). Cytochrome P450 9f2 genes have also been linked to insecticide resistance (Etebari et al., 2018). The four SNPs enclosing the region were detected as outliers for all 2 ≤ K ≤ 4 settings and were detected in both the imputed and unimputed datasets, which likely reflects the low missing data 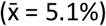 at these SNPs.

We investigated these four SNPs more closely using the unimputed dataset from the entire Indo-Pacific region (Table 3). Non-reference alleles at these SNPs were found most frequently as a single linked haplotype, which in the TSI was detected at highest frequency on I:Masig (2018 and 2019) but also on A:Keriri and L:Mer. Outside of the TSI this haplotype was found only in Port Moresby in PNG. Considering non-reference alleles at the four SNPs individually, outside of the TSI and Port Moresby the only observation was a single individual from Singapore, which had two alleles identical by state at one of the SNPs (Table 3). Tables S7 and S8 list unimputed genotypes at these SNPs for all TSI individuals (Table S7) and all non-TSI individuals (Table S8). These show clear but imperfect patterns of linkage between the two RADtags, and clear geographical restrictions in where the alleles were found.

**Table 3:** Frequencies of non-reference alleles (p) at the four outlier SNPs on scaffold NW_021838465.1. RADtag1 had one outlier SNP (pos: 84554160), RADtag2 had three outlier SNPs (pos: 84888984, 84888990, 84888991) which always cooccurred except in case of missing data. See Table S3 for a list of genes of known function within this region and Tables S7-S8 for genotypes of all individuals at the four outlier SNPs.

|  | p(84554160) | p(84888984) | p(84888990) | p(84888991) |
| --- | --- | --- | --- | --- |
| Location | RADtag1 | RADtag2 | RADtag2 | RADtag2 |
| **TSI** |  |  |  |  |
| A:Keriri | 0.056 | 0.045 | 0.045 | 0.045 |
| I:Masig (2018) | 0.238 | 0.272 | 0.272 | 0.272 |
| I:Masig (2019) | 0.250 | 0.250 | 0.250 | 0.250 |
| L:Mer | 0.040 | 0.021 | 0.021 | 0 |
| J:Ugar | 0 | 0.042 | 0.042 | 0.042 |
| All other locations | 0 | 0 | 0 | 0 |
| **International** |  |  |  |  |
| Port Moresby | 0.235 | 0.265 | 0.282 | 0.265 |
| Singapore | 0.071 | 0 | 0 | 0 |
| All other locations | 0 | 0 | 0 | 0 |

## Discussion

Spatial population genomics provides a powerful framework for investigating recent invasions. Our spatially consistent sampling within villages and simultaneous sampling across islands has permitted a range of analyses appropriate for characterising dispersal and adaptation in the recent *Ae. albopictus* invasion of the Torres Strait. Our findings point to a highly interconnected invasive metapopulation with frequent dispersal among both adjacent and non-adjacent islands, including incursions into uninvaded islands below the *cordon sanitaire*. The connectivity of this system also extends beyond borders, with evidence of gene flow from PNG into several of the nearest islands. Some of this gene flow was strongly suggestive of adaptive introgression, specifically at a set of linked outlier SNPs surrounding a genomic region containing 12 cytochrome P450 genes, of types (9e2 and 9f2) previously associated with insecticide resistance (Etebari et al., 2018; Kim Lien et al., 2019). Non-reference alleles at these outlier SNPs were found in four TSI villages, but not in any of the Indonesian populations from which the original invasion was sourced, or any population other than Port Moresby (PNG south coast), indicating a potential cross-country ‘genetic invasion’ of resistance alleles introgressed from southern PNG to the TSI. Broad-scale patterns of passive dispersal across islands contrasted with fine-scale patterns of active dispersal within villages, where spatial genetic structure was locally variable but displayed general trends of isolation by distance and neighbourhood size (N_W:_ Wright, 1946) of 232–577.

When geographical genetic structure is weak, as in the TSI (Fig 1, Fig 3), this is often read as a sign of high gene flow between subregions (Bossart & Prowell, 1998). However, in new invasions this conclusion can be confounded by high regional coancestry stemming from the original invasion (Cristescu, 2015; Fitzpatrick et al., 2012). This study instead investigated dispersal by analysing close kin dyads, incursions below the *cordon sanitaire*, and gene flow from PNG. Kinship and incursion tracing both reveal dispersal patterns over the past ∼1-2 generations, and these methods should not be confounded by regional coancestry as they were either calibrated against estimates from within villages (kinship) or traced incursive mosquitoes to anywhere in continuous space within the region (incursions: Locator). Of the four incursives, only Inc-4 was likely confounded by coancestry, as seen by its central placement among the training genotypes (Fig 2). Likewise, international gene flow had no trouble with coancestry as the source of gene flow (PNG) was genetically very distinct from the source of the initial invasion (Indonesia) (Fig 3, Figure S32). Together these analyses suggest that long-distance dispersal is frequent throughout the region, though movement between nearby islands may be more common than between distant islands (Fig 2, Table 2). Despite evidence of direct gene flow from PNG (Table 1), the TSI retains a distinct genetic background (Fig 3), indicating that if any local eradication has occurred as postulated in Muzari et al. (2017) these islands would have been recolonised primarily from within the TSI. The consistent clustering of PNG and Vanuatu genotypes indicate the Vanuatuan *Ae. albopictus* invasion was likely established from PNG or another location of that lineage like the Solomon Islands (see Maynard et al., 2017).

These patterns of variation are similar to those found in some biological invasions but contrast with others where variation reflects ongoing founder events as a colonising species spreads. Similar patterns to those observed here have been established for western flower thrips where high rates of ongoing gene flow mediated by human transport have been hypothesised (Cao et al., 2017). In contrast, regional genetic patterns reflective of repeated colonisations with small sample sizes have been documented in other thrips where distant colonisation into vacant habitat is likely to have occurred (Cao et al., 2019), as well as where there have been intensive campaigns with chemical insecticides to subdue populations (Cao et al., 2017). For TSI *Ae. albopictus*, it appears that human-mediated migration has had a stronger effect on population structure than control efforts aiming to suppress populations. In this sense, the TSI *Ae. albopictus* appear to be structured as a metapopulation similar to classical metapopulations like Glanville fritillary butterflies (Fountain et al., 2018; Hanski et al., 2017).

Although movement between islands has been suggested in previous studies (Beebe et al., 2013; Maynard et al., 2017), this study provides the first evidence of dispersal between specific islands. Our kinship analysis builds on previous uses of close kin to detect recent dispersal (M. Z. Chen et al., 2020; Escoda et al., 2019, 2017; Fountain et al., 2018; Jasper et al., 2019; Schmidt et al., 2018, 2017; Trense et al., 2020), and here demonstrates the use of regional genetic distance patterns to also infer the direction of movement (Fig 1, Table 2). Inferring movement patterns across single generations in this manner is becoming increasingly popular in applied work on pests and conservation, where recent changes in demography and genetic structure mean that assumptions of equilibrium demography are invalid and where anthropogenic impacts are observed from one generation to the next. First order kin detected across islands were likely transported via aeroplane, barge or small vessel movement. Although aircraft disinsection (treatment with insecticides) is currently compulsory for all air travel between the TSI region and the mainland of Australia, flights among the islands are not always disinsected. Future research could investigate the relative contributions of different transport networks to *Ae. albopictus* dispersal between islands and into the TSI from PNG.

The power of higher-resolution markers to better delineate geographical patterns among genotypes is well-established (Escoda et al., 2017; Puckett & Eggert, 2016; Rašić et al., 2014). While previous investigation of this system with lower-resolution markers detected some spatial structuring among islands (Maynard et al., 2017), this study found no greater sorting of genotypes by location despite higher marker power and despite sampling in this study having taken place 3-11 years later (Fig 1, Fig 3). These results point to the TSI *Ae. albopictus* population remaining highly interconnected over time. This interconnection is likely the reason for high genetic variability within villages, including the inconsistent spatial structuring of genetic variation among dyads (Table 1). Nevertheless, our estimate of N_W_ in villages (232 – 577) is consistent with a previous estimate for an urban *Ae. aegypti* population (268; Jasper et al., 2019). Considering high densities of *Ae. albopictus* in the TSI, this estimate of N_W_ is suggestive of some spatial structure within villages.

Despite high regional gene flow, we managed to locate the source of 2-3 of the 4 incursions below the *cordon sanitaire*. Considering the absence of discrete populations within the TSI, the genotype-based method of Locator (Battey et al., 2020) was better suited to this dataset than the allele-frequency based method of assignPOP (K.-Y. Chen et al., 2018), though the latter still performed well for Inc-1 and Inc-2 and both methods successfully traced samples collected 10 months after the initial collections. Although Locator may best be suited to non-clustered sampling designs that are rare in island systems, Locator may sidestep issues relating to invasion age and frequent gene flow by not requiring that reference genotypes be sorted into populations. Locator’s placement of Inc-4 across a wide swathe of the sampling range (Fig 2) indicated that this sample could not be assigned, which was not likely due to missing data but due to recent ancestry from multiple locations. If we treat the placement of Inc-4 as the expected outcome of a failure to assign, then the placement of Inc-3 on Moa Island with closely-clustered bootstrap replicates suggests that this is its likely origin. While population assignment methods such as assignPOP perform well when determining incursion sources pre-invasion (Schmidt, Chung, van Rooyen, et al., 2020; Schmidt et al., 2019), Locator appears a more useful tool for tracing local incursions after an initial invasion.

Detecting adaptive introgression is a powerful application of genomic datasets. For invasive systems adaptive introgression can be classed as a type of genetic invasion, in which alleles conferring a selective advantage that may interfere with control strategies are introduced after an initial invasion (Endersby-Harshman et al., 2020; Riveron et al., 2013; Schmidt et al., 2019). Our detection of a set of alleles from PNG that have introgressed into the TSI is strongly suggestive of a genetic invasion. In the TSI, introgressed alleles at the four outlier SNPs were found on four of the islands, but were particularly common on I:Masig, where their frequency remained stable between 2018 and 2019. Considering the current low allele frequencies on most of the islands, the imperfect linkage between the alleles, and the abandonment of the insecticide program in the TSI a decade previously (Muzari et al., 2017), our observations may reflect the aftermath of a genetic invasion that spread into the TSI during the height of the insecticide program. During this time, I:Masig may have been acting as an ‘invasive bridgehead’ (Estoup & Guillemaud, 2010) for the genetic invasion, from which resistance alleles spread to other islands.

While traditionally ‘introgression’ refers to the interspecific transfer of genetic material, which has also been observed in mosquitoes (Norris et al., 2015), introgression between differentiated populations as observed here may be more common and may allow for faster spread of advantageous alleles if interspecific mating is rare or confers greater fitness costs. Considering also the connectivity of the TSI/PNG system, where dispersal takes place among distant islands and across countries, if advantageous alleles arise in any given location they may rapidly spread throughout the region if similar selective pressures are present. This rapid spread of alleles may be a threat in any biological system with high dispersal rates and common selective pressures. In the TSI, linkage was strong between the two RADtags surrounding the 12 cytochrome P450 genes, though not as strong as the near-perfect linkage observed around pyrethroid resistance mutations in *Ae. aegypti* (Endersby-Harshman et al., 2020). This may reflect relatively weaker selection around the cytochrome P450 genes given that area-wide insecticide use in the TSI was phased out ∼10 years before sampling for this project (Muzari et al., 2017) and that in PNG most documented insecticide use occurs outside of cities like Port Moresby (Demok et al., 2019). While recent bioassays on PNG *Ae. albopictus* demonstrated resistance to DDT, these investigated only Madang *Ae. albopictus* (Demok et al., 2019), though emerging results indicate these resistance patterns are similar across PNG (unpublished data). Nevertheless, the detected introgression was evidently not from Madang but from either Port Moresby or an unsampled location in PNG. Future work is needed to determine what role the 12 cytochrome P450 genes have in conferring insecticide resistance in this system and in *Ae. albopictus* more broadly.

## Conclusions

Dispersal can be difficult to investigate in many biological systems, particularly those subjected to different ongoing dispersal processes. The recent biological invasion of *Ae. albopictus* into TSI provides an example of such a system, and here we have performed a comprehensive characterisation of dispersal using temporally-restricted sampling and an appropriate set of population genomic analyses. We detected recent dispersal between distant and adjacent islands and found no increase in spatial genetic structure over time, suggesting that the invaded islands may best be considered a metapopulation connected by frequent gene flow, rather than individual ‘populations’. We also made use of the strongly-differentiated PNG *Ae. albopictus* population to detect gene flow from PNG into specific islands, and found evidence that a genetic invasion of insecticide resistance alleles may have spread into the TSI from the south coast of PNG. These methods will be broadly applicable to other taxa, particularly those affected by anthropogenic processes. For *Ae. albopictus*, the findings inform strategies for the control of this globally-invasive pest across a broad range of spatial scales.

## Acknowledgements

For sample collection we thank Esther Anderson, Moses Laman, Rotarians against Malaria PNG staff, and the PNGIMR Vector Borne Disease Unit entomology team. We thank Moshe Jasper for helpful discussions. For funding we thank Queensland Health, the Commonwealth Department of Health, the Far North Queensland Hospital Foundation (FNQHF), and the NHMRC (Programme Grant 1037003, Career Development Fellowship 1141441, and Fellowship Grants 1118640 and 5121190). T.S. was financially supported by a Australian Government Research Training Program Scholarship.

## Data Accessibility

Aligned .bam sequence files for 331 TSI *Ae. albopictus*, 4 incursive *Ae. albopictus*, and 36 PNG *Ae. albopictus* are available through the Sequence Read Archive at NCBI Genbank: PRJNA684450.

## Author Contributions

TLS, TS, GE, SR, and AAH designed the study. TS, SK, SD, QY, MOM, GE, MB, RB and PH collected the data. TLS, TS, JC and MAF analysed the data. TLS, MAF, GE, TRB, SR and AAH provided supervision. TLS, TS and AAH wrote the paper with edits from all authors.

## Supporting Information Legends

### Text

Text S1: Details of the genotyping pipeline.

### Tables

Table S1: Initial assignPOP run showing assignment probability to each TSI and non-TSI location, and aggregated assignment probabilities for all TSI and all non-TSI locations.

Table S2: Pairwise FST estimates among TSI villages.

Table S3: The first of four genomic regions containing multiple outlier SNPs found on multiple RADtags, and the genes of known product or function located between these RADtags.

Table S4: The second of four genomic regions containing multiple outlier SNPs found on multiple RADtags, and the genes of known product or function located between these RADtags.

Table S5: The third of four genomic regions containing multiple outlier SNPs found on multiple RADtags, and the genes of known product or function located between these RADtags.

Table S6: The fourth of four genomic regions containing multiple outlier SNPs found on multiple RADtags, and the genes of known product or function located between these RADtags.

Table S7: Non-reference alleles at the four outlier SNPs in Region 1 (see Table S2) found in TSI villages.

Table S8: Non-reference alleles at the four outlier SNPs in Region 1 (see Table S2) found in non-TSI locations.

### Figures

Figure S1: Locations of *Aedes albopictus* sweep net collections on Badu Island. Map produced using Mapinfo (2019) with the Queensland basemap satellite imagery (2020).

Figure S2: Locations of *Aedes albopictus* sweep net collections on Dauan Island. Map produced using Mapinfo (2019) with the Queensland basemap satellite imagery (2020).

Figure S3: Locations of *Aedes albopictus* sweep net collections on Erub Island. Map produced using Mapinfo (2019) with the Queensland basemap satellite imagery (2020).

Figure S4: Locations of *Aedes albopictus* sweep net collections on Iama Island. Map produced using Mapinfo (2019) with the Queensland basemap satellite imagery (2020).

Figure S5: Locations of *Aedes albopictus* sweep net collections on Keriri Island. Map produced using Mapinfo (2019) with the Queensland basemap satellite imagery (2020).

Figure S6: Locations of *Aedes albopictus* sweep net collections on Mabuiag Island. Map produced using Mapinfo (2019) with the Queensland basemap satellite imagery (2020).

Figure S7: Locations of *Aedes albopictus* sweep net collections on Masig Island. Map produced using Mapinfo (2019) with the Queensland basemap satellite imagery (2020).

Figure S8: Locations of *Aedes albopictus* sweep net collections on Poruma Island. Map produced using Mapinfo (2019) with the Queensland basemap satellite imagery (2020).

Figure S9: Locations of *Aedes albopictus* sweep net collections on Mer Island. Map produced using Mapinfo (2019) with the Queensland basemap satellite imagery (2020).

Figure S10: Locations of *Aedes albopictus* sweep net collections on Ugar Island. Map produced using Mapinfo (2019) with the Queensland basemap satellite imagery (2020).

Figure S11: Locations of *Aedes albopictus* sweep net collections on St Pauls (Moa Island). Map produced using Mapinfo (2019) with the Queensland basemap satellite imagery (2020).

Figure S12: Locations of *Aedes albopictus* sweep net collections on Kubin (Moa Island). Map produced using Mapinfo (2019) with the Queensland basemap satellite imagery (2020).

Figure S13: Locations of *Aedes albopictus* sweep net collections on Warraber Island. Map produced using Mapinfo (2019) with the Queensland basemap satellite imagery (2020).

Figure S14: Location of sweep net sampling sites and detected incursions on Ngurapai between 2019-02-02 and 2019-02-09. Map produced using Mapinfo (2019) with the Queensland basemap satellite imagery (2020).

Figure S15: Individual ancestral lineage proportions for each village (n=22) with two ancestral lineages (K = 2) selected.

Figure S16: Individual ancestral lineage proportions for each village (n=22) with three ancestral lineages (K = 3) selected.

Figure S17: Individual ancestral lineage proportions for each village (n=22) with four ancestral lineages (K = 4) selected.

Figure S18: Scatter plot of a mantel test between geographic (natural log) and genetic distance (Rousset’s a) for *Aedes albopictus* collections on Badu Island. Lines describe a linear regression fit with 95% Confidence Interval shown.

Figure S19: Scatter plot of a mantel test between geographic (natural log) and genetic distance (Rousset’s a) for *Aedes albopictus* collections on Keriri Island. Lines describe a linear regression fit with 95% Confidence Interval shown.

Figure S20: Scatter plot of a mantel test between geographic (natural log) and genetic distance (Rousset’s a) for *Aedes albopictus* collections on Masig Island in 2018. Lines describe a linear regression fit with 95% Confidence Interval shown.

Figure S21: Scatter plot of a mantel test between geographic (natural log) and genetic distance (Rousset’s a) for *Aedes albopictus* collections on Masig Island in 2019. Lines describe a linear regression fit with 95% Confidence Interval shown.

Figure S22: Scatter plot of a mantel test between geographic (natural log) and genetic distance (Rousset’s a) for *Aedes albopictus* collections on Mer Island. Lines describe a linear regression fit with 95% Confidence Interval shown.

Figure S23: Scatter plot of a mantel test between geographic (natural log) and genetic distance (Rousset’s a) for *Aedes albopictus* collections on Warraber Island. Lines describe a linear regression fit with 95% Confidence Interval shown.

Figure S24: Scatter plot of a mantel test between geographic (natural log) and genetic distance (Rousset’s a) for *Aedes albopictus* collections on Erub Island. Lines describe a linear regression fit with 95% Confidence Interval shown.

Figure S25: Scatter plot of a mantel test between geographic (natural log) and genetic distance (Rousset’s a) for *Aedes albopictus* collections on Mabuiag Island. Lines describe a linear regression fit with 95% Confidence Interval shown.

Figure S26: Scatter plot of a mantel test between geographic (natural log) and genetic distance (Rousset’s a) for *Aedes albopictus* collections on Poruma Island. Lines describe a linear regression fit with 95% Confidence Interval shown.

Figure S27: Scatter plot of a mantel test between geographic (natural log) and genetic distance (Rousset’s a) for *Aedes albopictus* collections on Iama Island. Lines describe a linear regression fit with 95% Confidence Interval shown.

Figure S28: Scatter plot of a mantel test between geographic (natural log) and genetic distance (Rousset’s a) for *Aedes albopictus* collections on Ugar Island. Lines describe a linear regression fit with 95% Confidence Interval shown.

Figure S29: Scatter plot of a mantel test between geographic (natural log) and genetic distance (Rousset’s a) for *Aedes albopictus* collections on Kubin (Moa Island). Lines describe a linear regression fit with 95% Confidence Interval shown.

Figure S30: Scatter plot of a mantel test between geographic (natural log) and genetic distance (Rousset’s a) for *Aedes albopictus* collections on St Pauls (Moa Island). Lines describe a linear regression fit with 95% Confidence Interval shown.

Figure S31: Full-size figure of Fig 3.

Figure S32: PCA-UMAP of all non-incursive genotypes, using the first 4 PCs, 50 neighbours and a 0.5 minimum distance.

Figure S33: Scree plot from pcadapt.

Figure S34: Genome-wide genetic structure of TSI villages, for the first and second PCs.

Figure S35: Genome-wide genetic structure of TSI villages, for the third and fourth PCs.

Figure S36: Genome-wide genetic structure of TSI villages, for the fifth and sixth PCs.

Figure S37: Genome-wide genetic structure of TSI villages, for the seventh and eighth PCs.

