## Supplementary material for "Spatial population genomics of a recent mosquito invasion": Figures S1-S37, Text S1, Tables S1-S2

* equal first authors

^ correspondence

[Figure S8:	1](#_Toc49242368)1

[Figure S9:	1](#_Toc49242369)2

[Figure S10:	1](#_Toc49242370)3

[Figure S11:	1](#_Toc49242371)4

[Figure S12:	1](#_Toc49242372)5

[Figure S13:	1](#_Toc49242373)6

[Figure S14	1](#_Toc49242374)7

[References for Figures S1-14	1](#_Toc49242374)8

[Text S1	1](#_Toc49242374)9

[Figure S15	2](#_Toc49242374)1

[Figure S23:](#_Toc49242366)	29

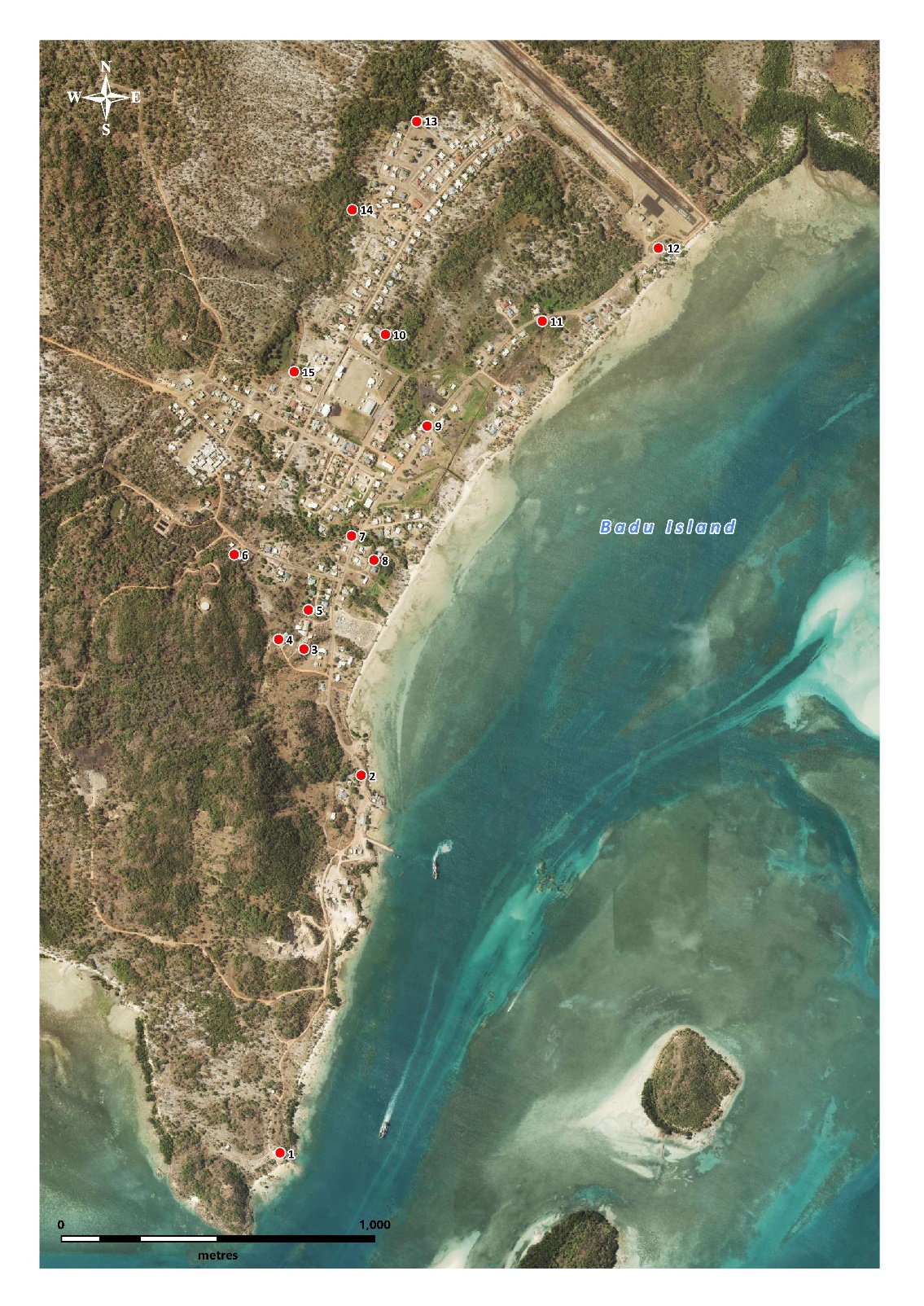

Figure S1: Locations of *Aedes albopictus* sweep net collections on Badu Island. Map produced using Mapinfo (2019) with the Queensland basemap satellite imagery (2020).

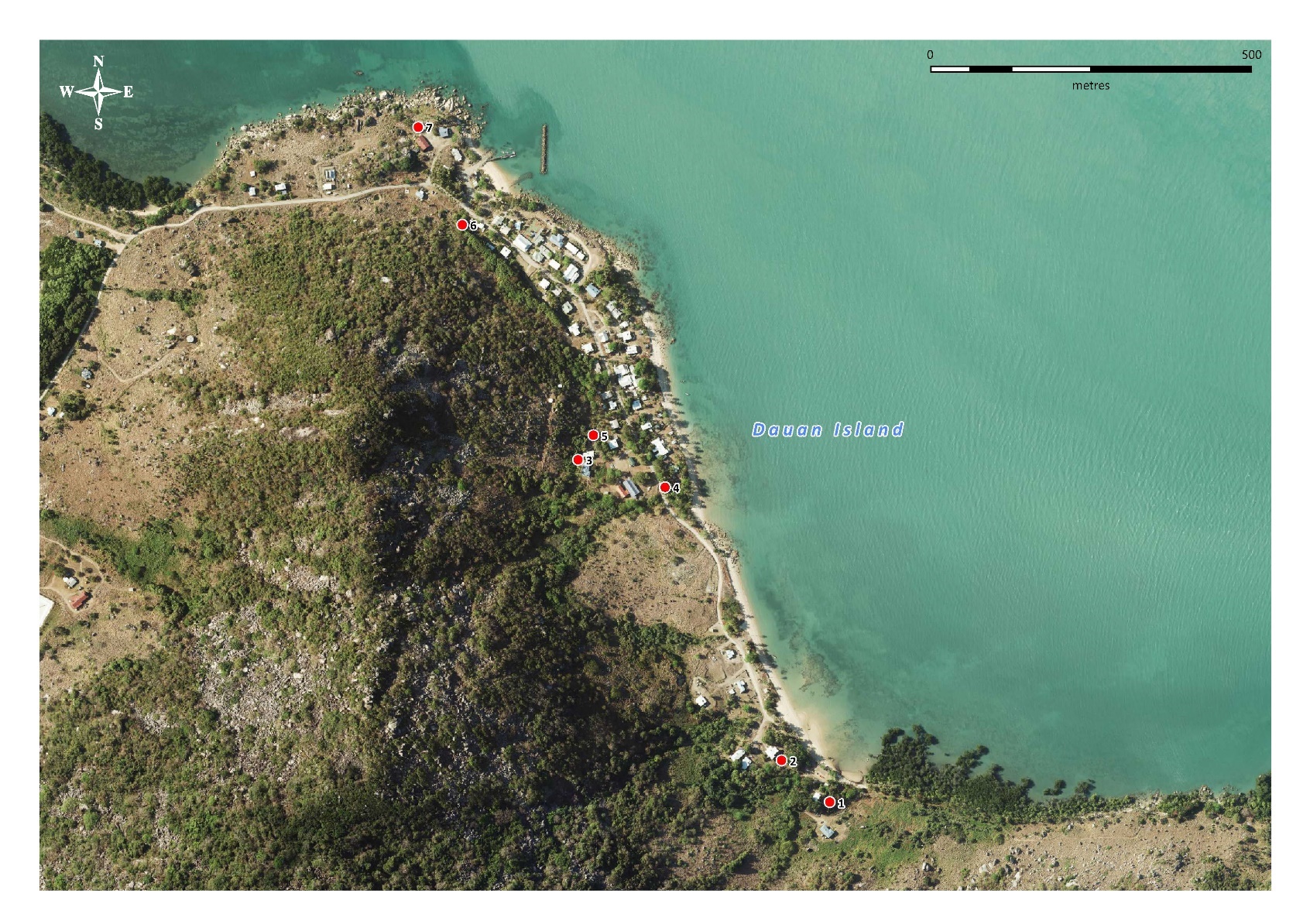

Figure S2: Locations of *Aedes albopictus* sweep net collections on Dauan Island. Map produced using Mapinfo (2019) with the Queensland basemap satellite imagery (2020).

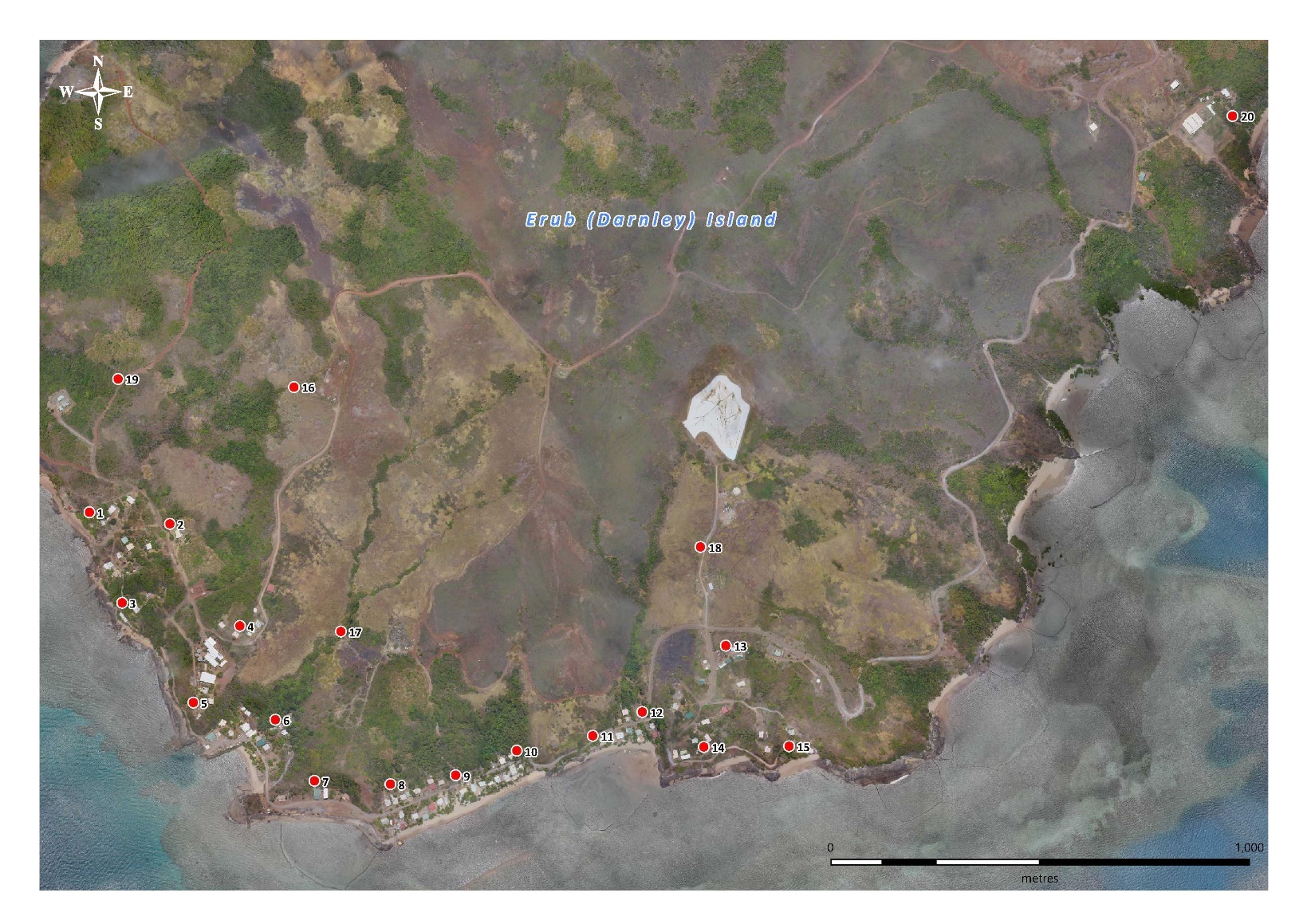

Figure S3: Locations of *Aedes albopictus* sweep net collections on Erub Island. Map produced using Mapinfo (2019) with the Queensland basemap satellite imagery (2020).

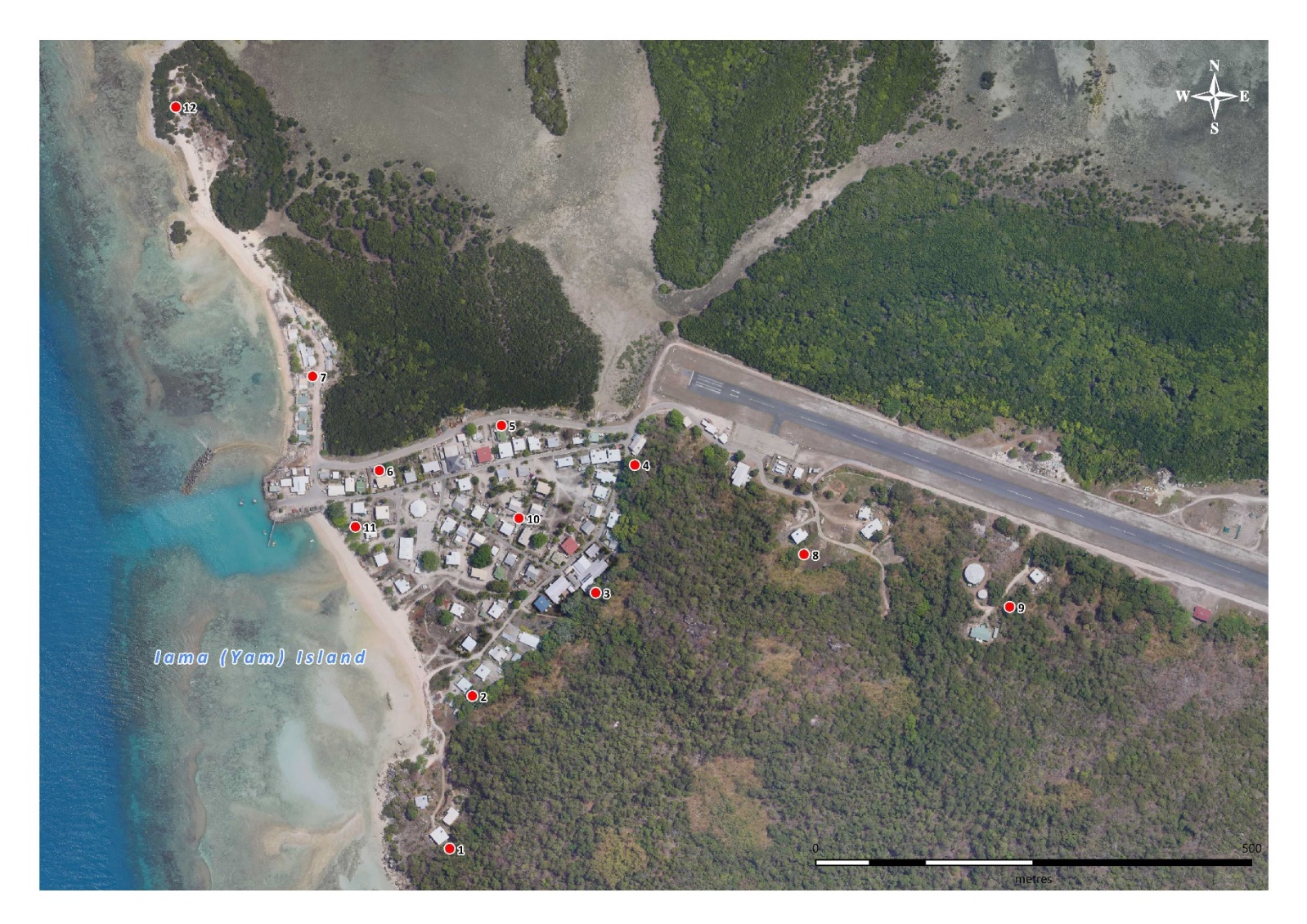

Figure S4: Locations of *Aedes albopictus* sweep net collections on Iama Island. Map produced using Mapinfo (2019) with the Queensland basemap satellite imagery (2020).

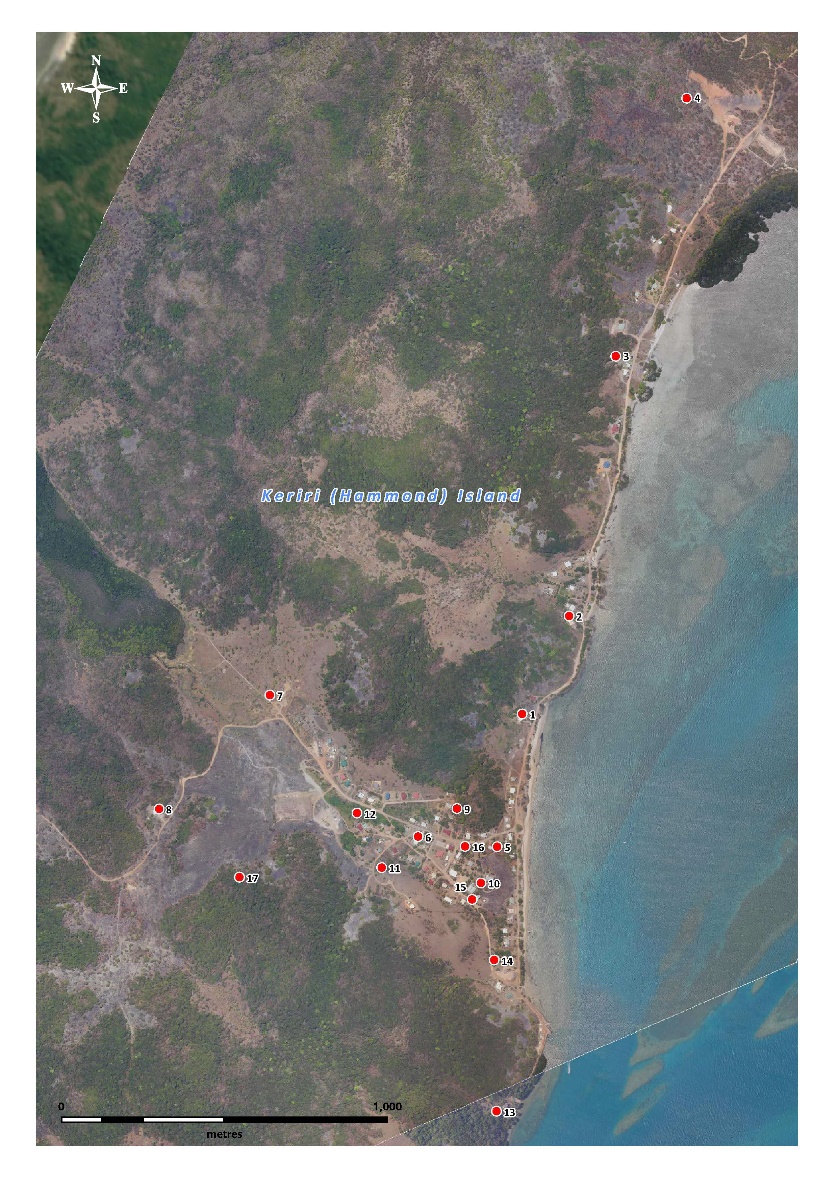

Figure S5: Locations of Aedes albopictus sweep net collections on Keriri Island. Map produced using Mapinfo (2019) with the Queensland basemap satellite imagery (2020).

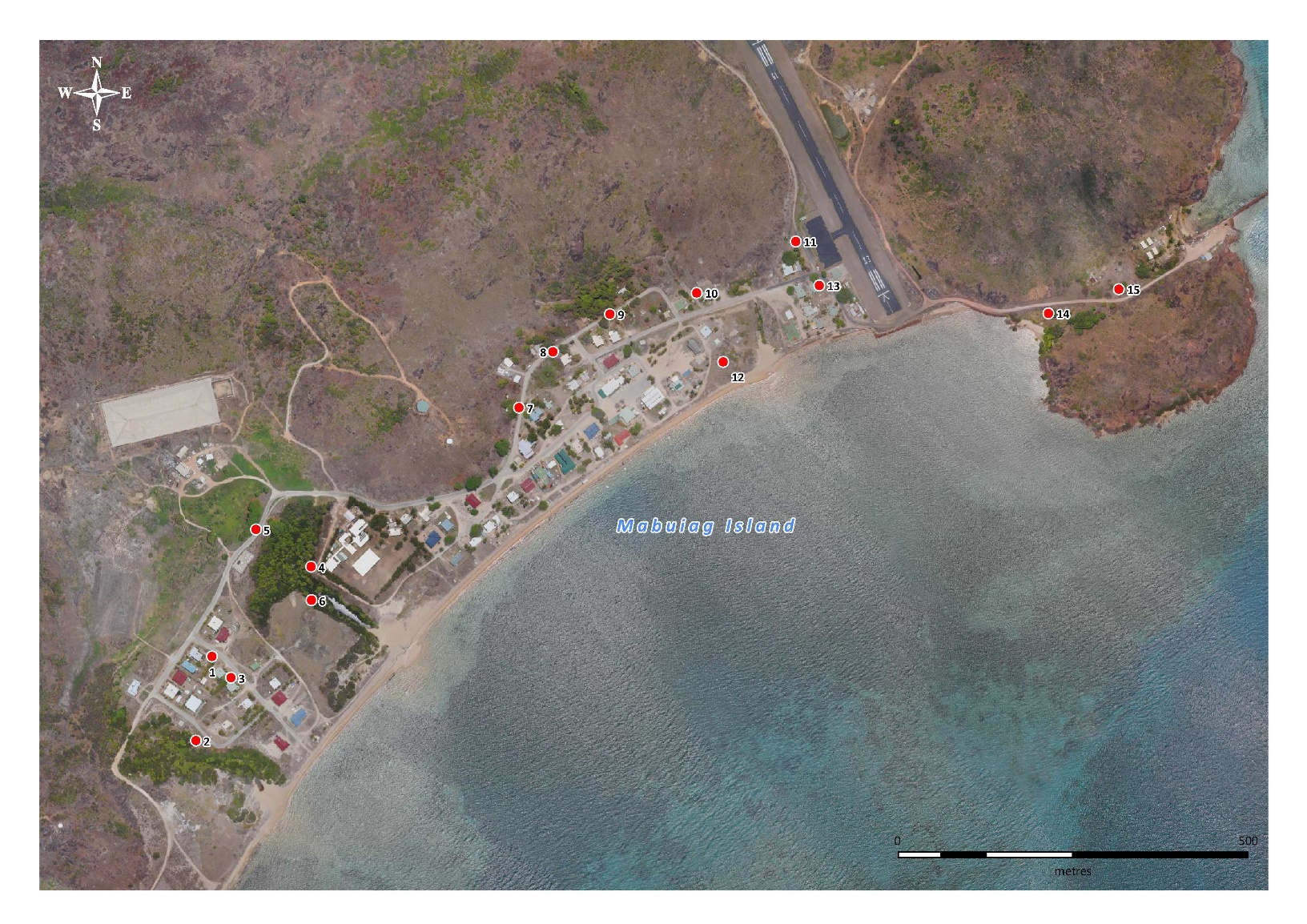

Figure S6: Locations of Aedes albopictus sweep net collections on Mabuiag Island. Map produced using Mapinfo (2019) with the Queensland basemap satellite imagery (2020).

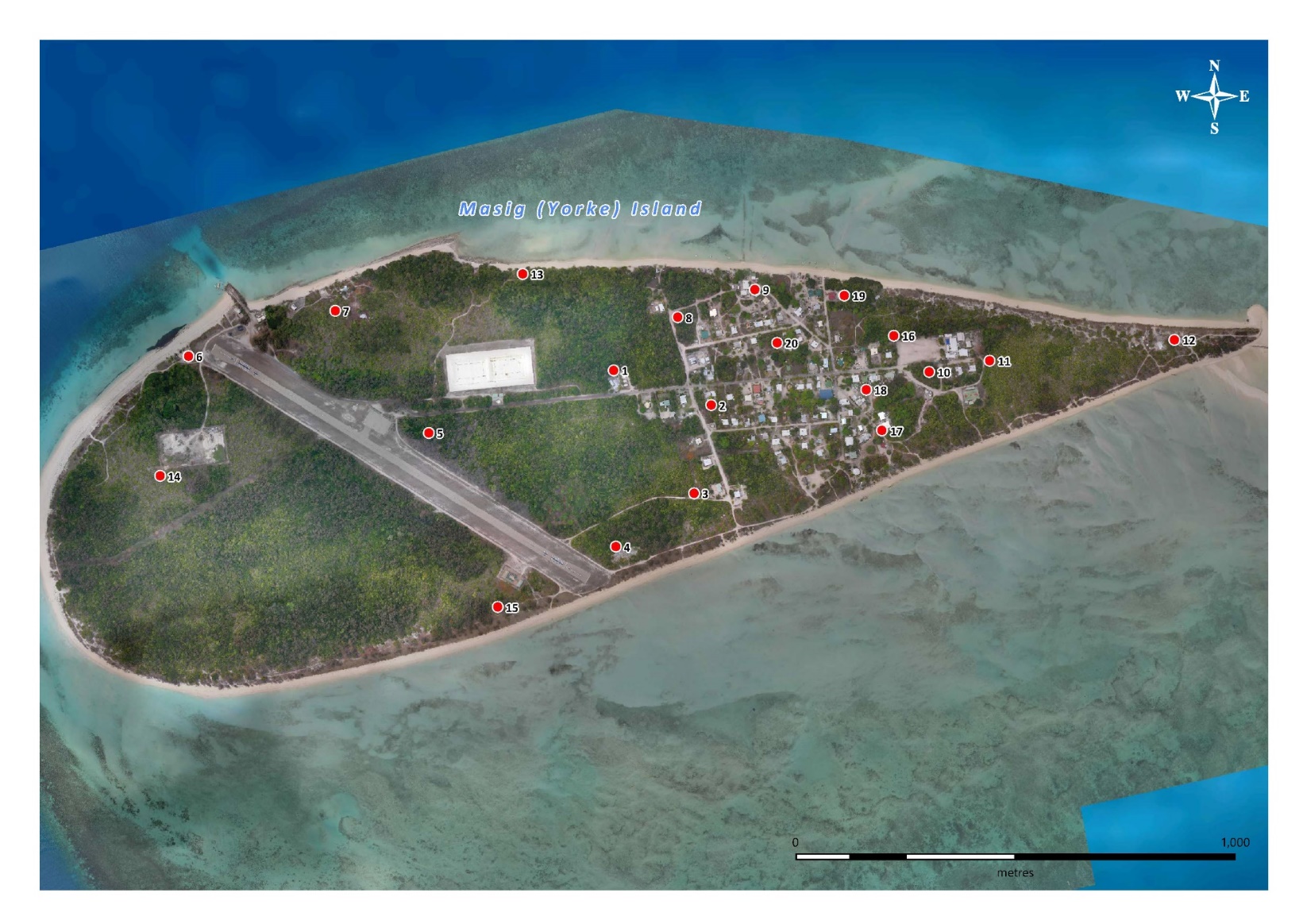

Figure S7: Locations of Aedes albopictus sweep net collections on Masig Island. Map produced using Mapinfo (2019) with the Queensland basemap satellite imagery (2020).

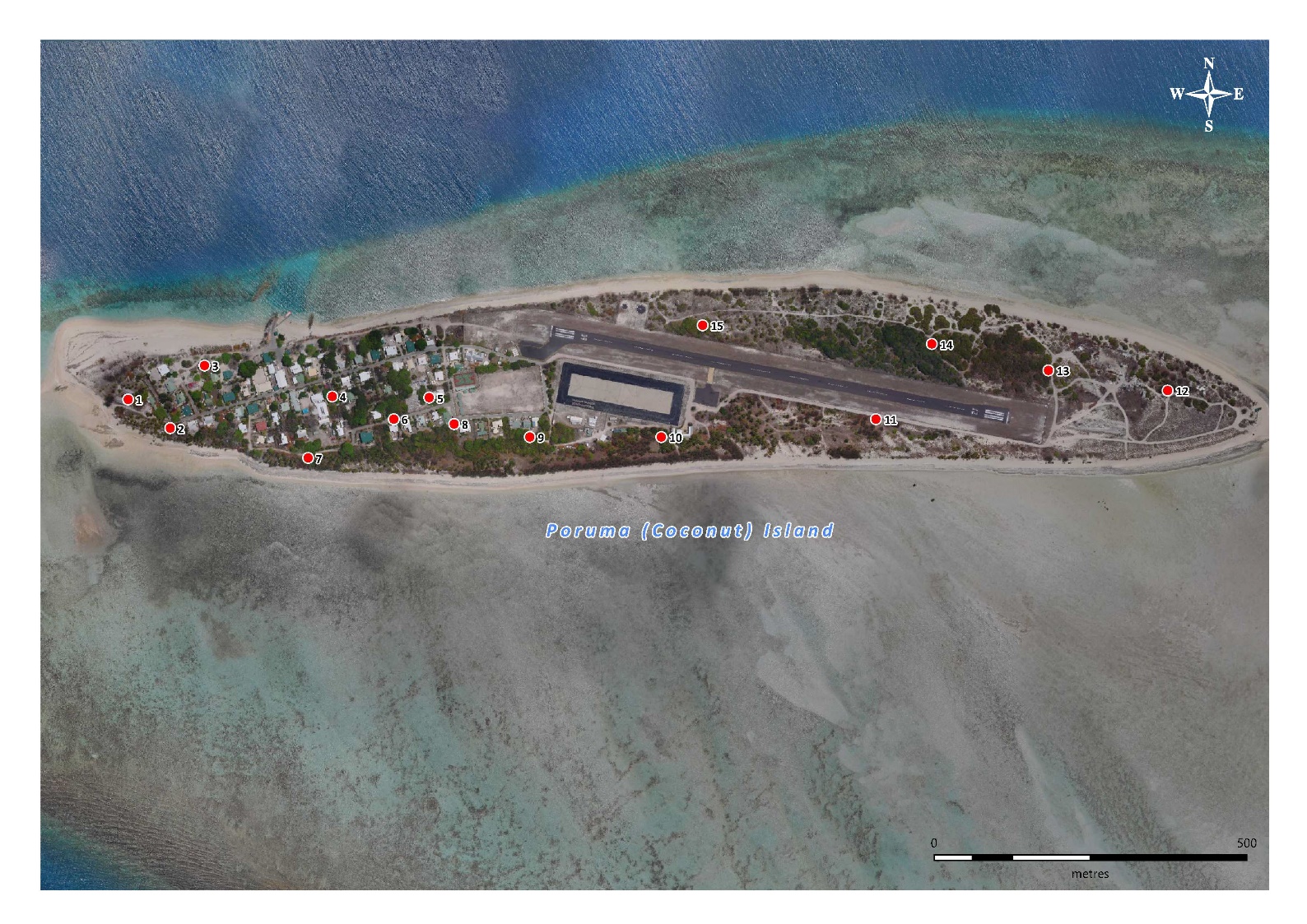

Figure S8: Locations of *Aedes albopictus* sweep net collections on Poruma Island. Map produced using Mapinfo (2019) with the Queensland basemap satellite imagery (2020).

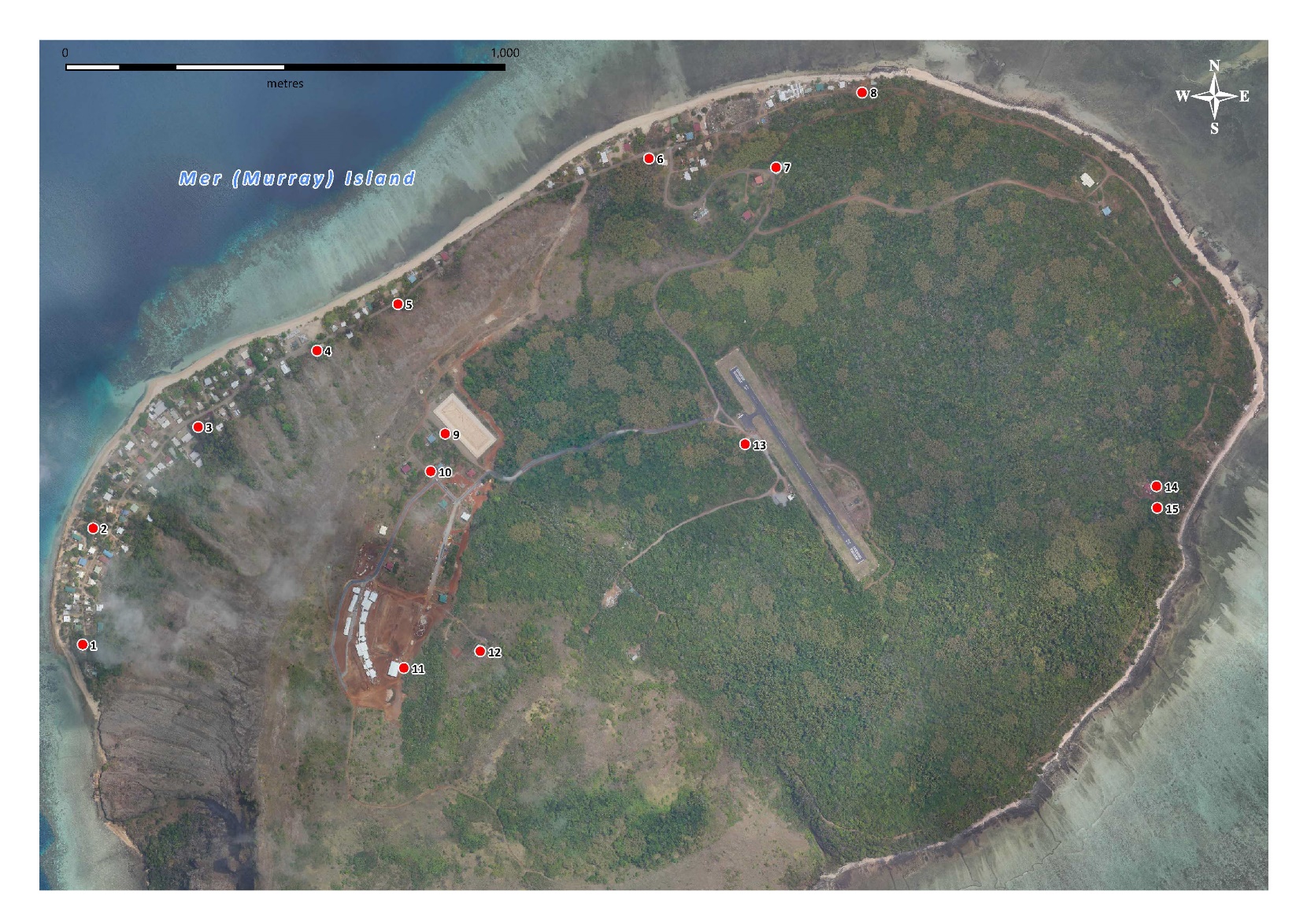

Figure S9: Locations of *Aedes albopictus* sweep net collections on Mer Island. Map produced using Mapinfo (2019) with the Queensland basemap satellite imagery (2020).

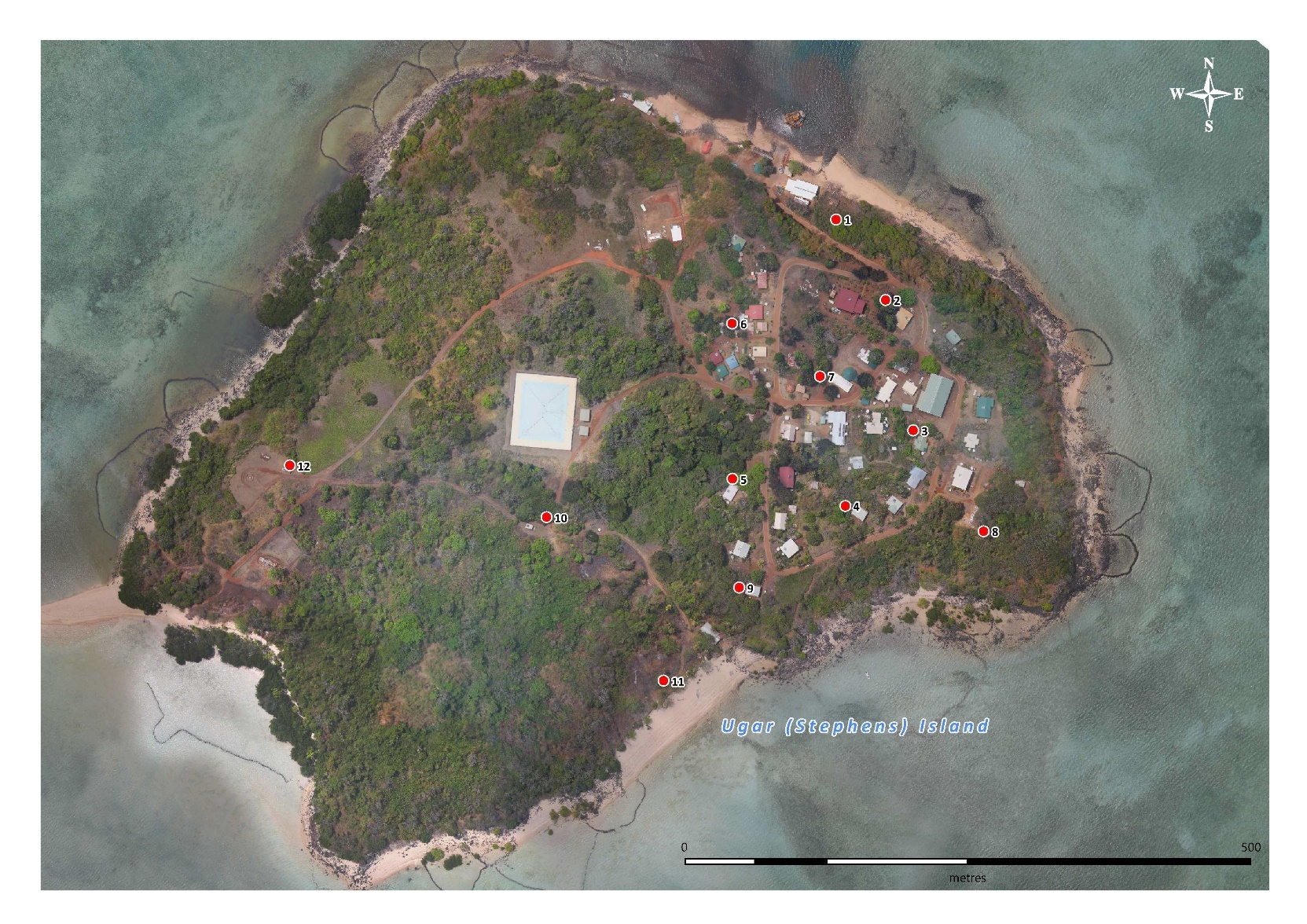

Figure S10: Locations of *Aedes albopictus* sweep net collections on Ugar Island. Map produced using Mapinfo (2019) with the Queensland basemap satellite imagery (2020).

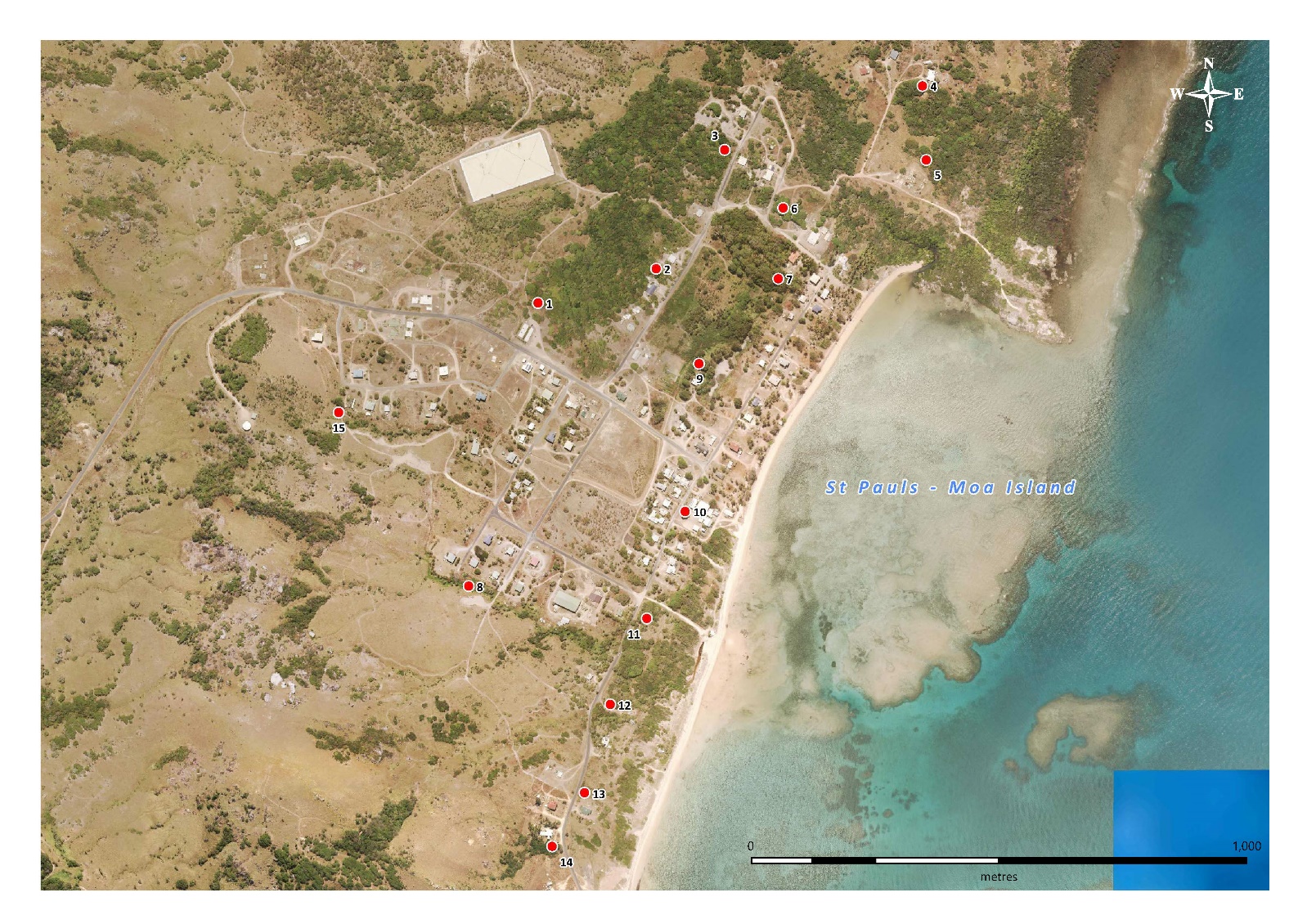

Figure S11: Locations of *Aedes albopictus* sweep net collections on St Pauls (Moa Island). Map produced using Mapinfo (2019) with the Queensland basemap satellite imagery (2020).

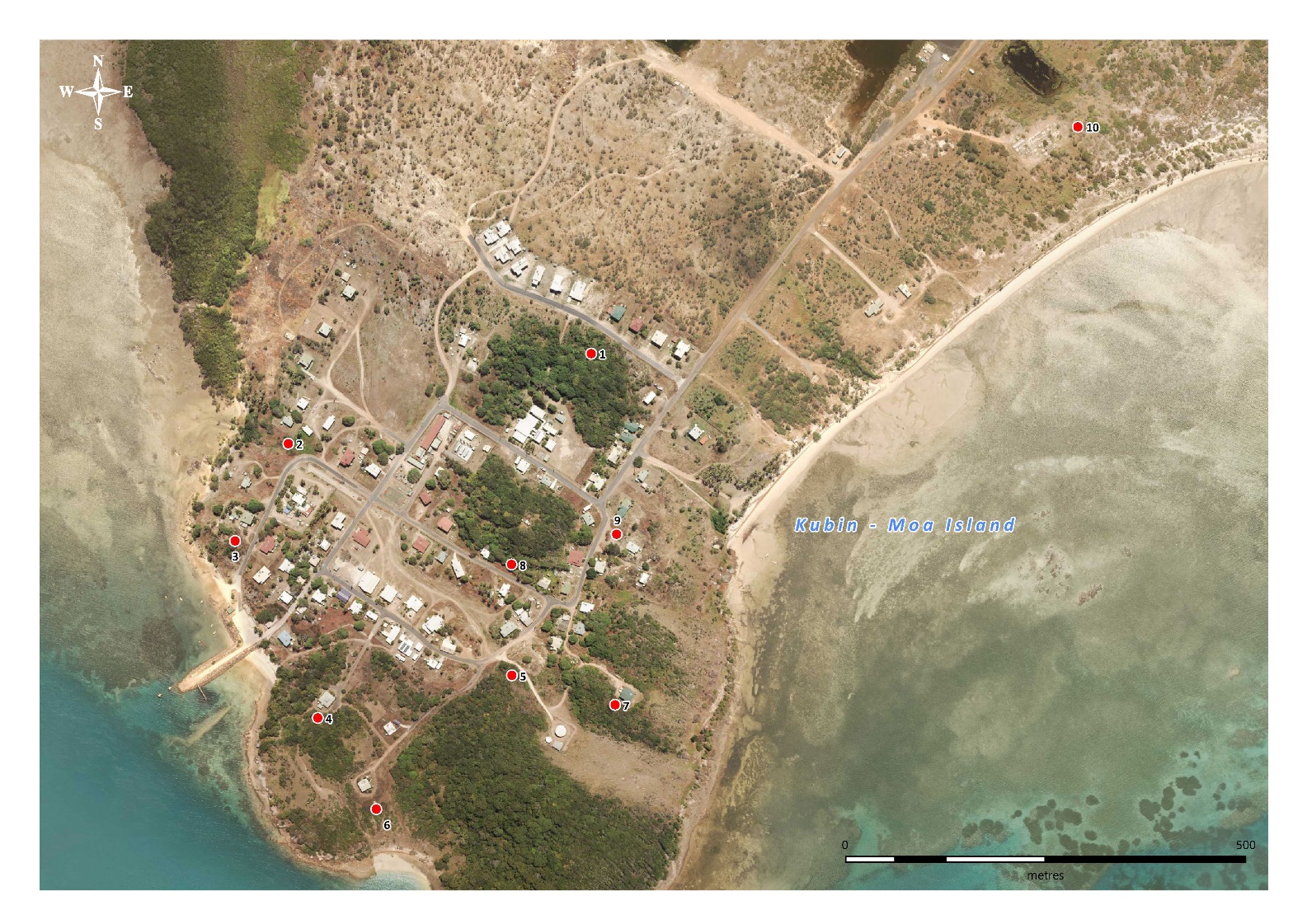

Figure S12: Locations of *Aedes albopictus* sweep net collections on Kubin (Moa Island). Map produced using Mapinfo (2019) with the Queensland basemap satellite imagery (2020).

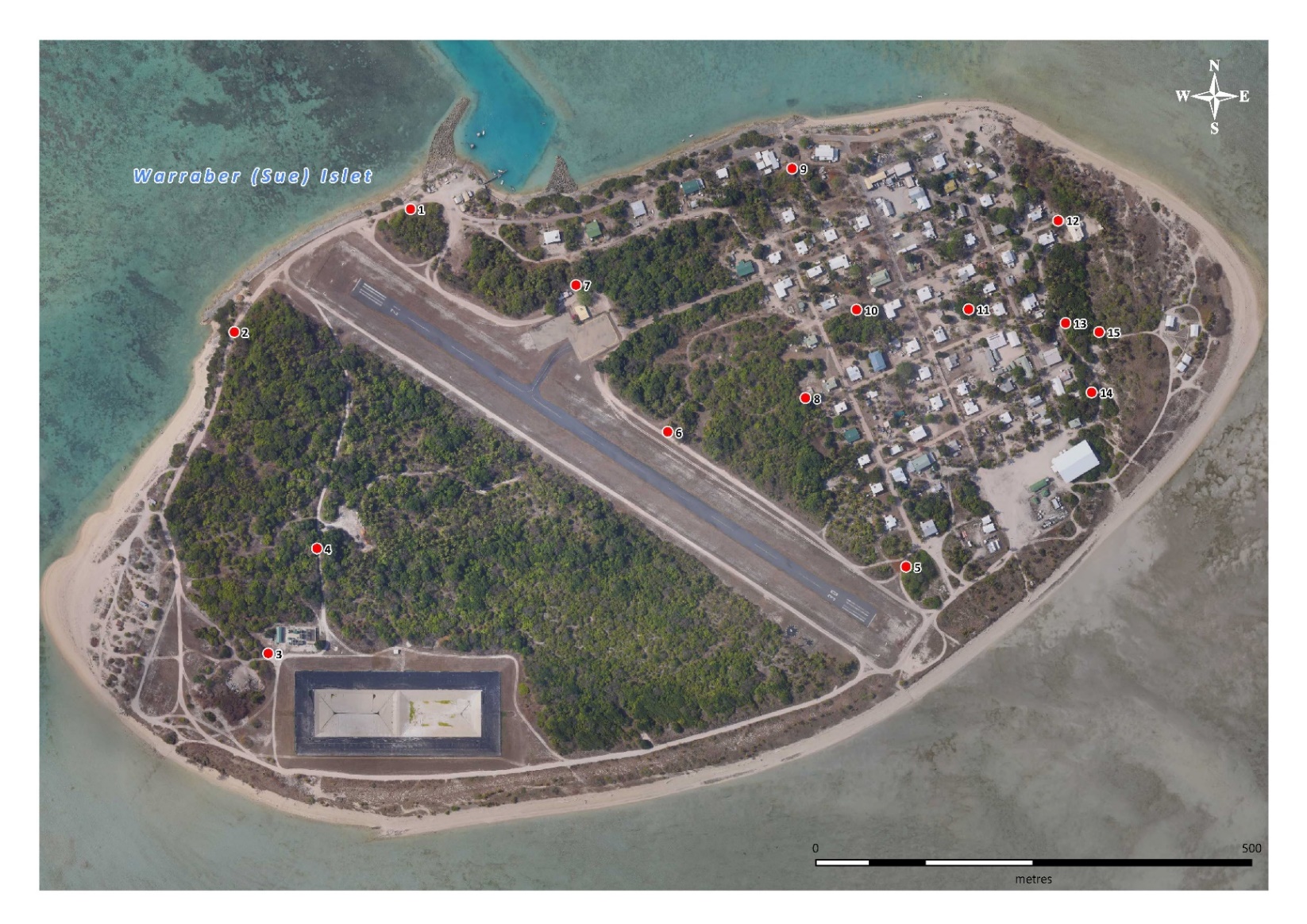

Figure S13: Locations of *Aedes albopictus* sweep net collections on Warraber Island. Map produced using Mapinfo (2019) with the Queensland basemap satellite imagery (2020).

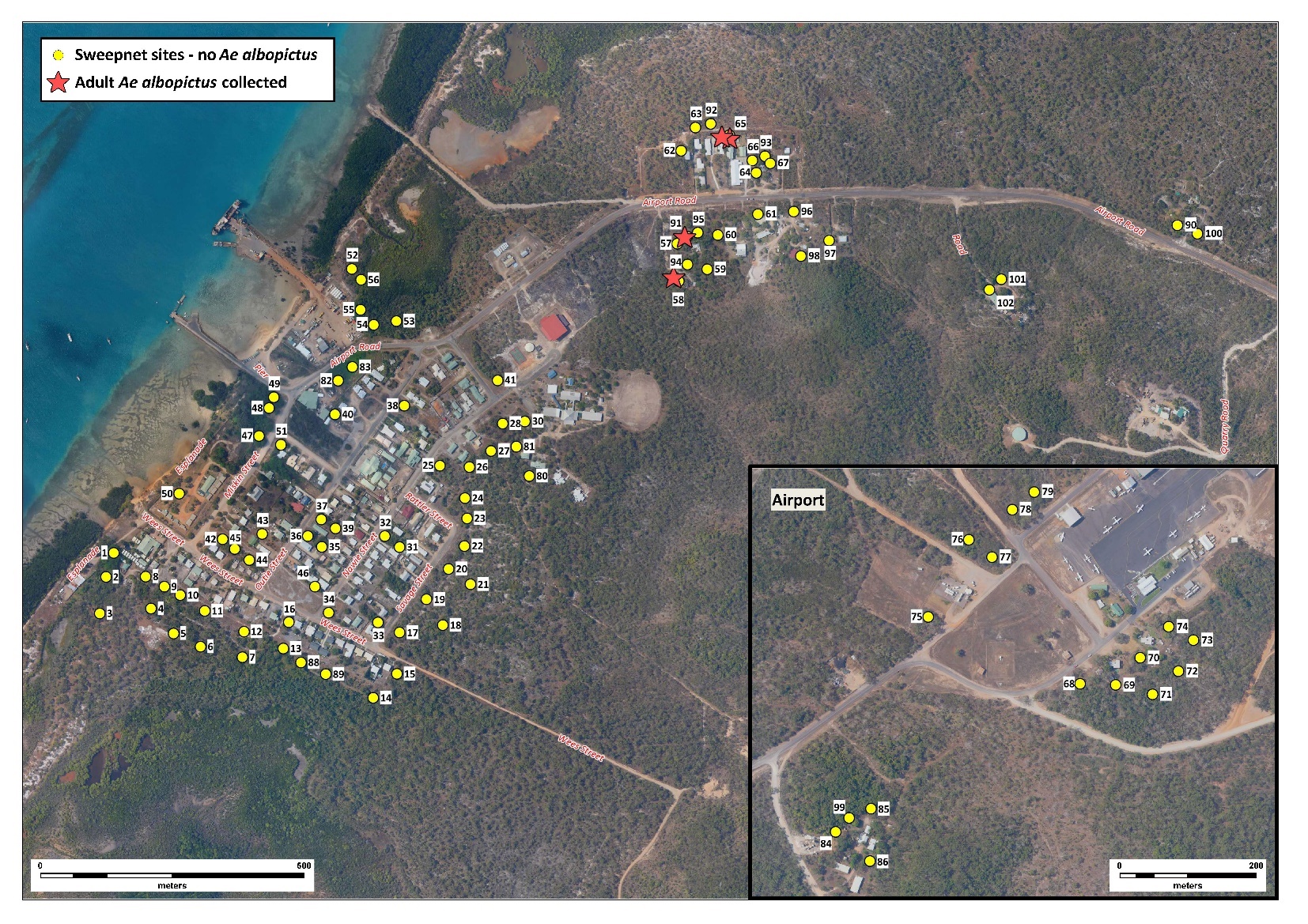

Figure S14: Location of sweep net sampling sites and detected incursions on Ngurapai between 2019-02-02 and 2019-02-09. Map produced using Mapinfo (2019) with the Queensland basemap satellite imagery (2020).

### Supplementary Text S1:

***Experimental design: laboratory procedures and sample processing***

Inadequate experimental design at the laboratory phase can produce confounding effects (i.e. large batch effect), capable of greatly biasing results (Bálint et al. 2018). As such, the following measures were taken: i) the investigator was blinded to the sample identity in the laboratory by using a unique set of identification codes, ii) samples were randomly selected on the day of DNA extraction and iii) the six libraries created contained on average the same number of randomly selected individuals from each island (60 – 64 individuals per library).

***DNA extraction & library preparation***

Mosquitoes were morphologically identified using keys from Webb et al. (2016) and sexed prior to DNA extraction. DNA extraction was conducted following manufacturer’s instructions using either a DNeasy Blood & Tissue kit (Qiagen, Hilden, Germany) or a Roche High Pure PCR Template Preparation Kit (Roche Molecular Systems, Inc., Pleasanton, CA, USA), with an additional RNAse A step. Resulting DNA extractions were stored at -80°C until selected for library preparation.

Double‐digest restriction site‐associated DNA sequencing (ddRADseq) libraries were prepared using a protocol developed by Rašić et al. (2014) for *Ae. aegypti* and validated in *Ae. albopictus* by Schmidt et al. (2017). Individuals from different sampling locations were selected randomly for inclusion in libraries.

120 ng of DNA was aliquoted from each library in a 45 uL reaction with 10 units of NlaIII and MluCI restriction enzymes (New England Biolabs, Beverly MA, USA), NEB CutSmart® buffer, and water. Digestions were incubated for 3.5 hours at 37°C with no heat kill step. 60 uL of Sera-Mag beads (1.5x concentration) were used to clean the digested products and this was ligated to unique Illumina P1 and P2 adapters with 1,000 units of T4 ligase (New England Biolabs, Beverly, MA, USA) incubated at 16°C overnight, with a final heat deactivation step at 65°C for 10 minutes. Our combinatorial indexing system comprising of 16 P1 adapters and 4 P2 adapters allowed for unique DNA barcodes to be ligated to each individual. Ligation products were pooled into a single 1.7 mL Eppendorf™ tube and cleaned with 1.5x concentration of Sera-Mag beads.

Size selection retaining DNA fragments between 300-450 base pairs (bp) was achieved using a Pippin-Prep 2% gel cassette (Sage Sciences, Beverly, MA, USA). Eight separate 1 μL volumes of size‐selected DNA from each libraries were used as templates in a 10 μL PCR reaction with 2.7 μL of water, 2 μL Phusion 5x buffer, 0.2 μL of dNTPs (10 mM), 2 μL of Illumina primer F (10 μM), 2 μL of Illumina primer R (10 μM) and 0.10 μL of Phusion Polymerase. PCR conditions were: 98°C for 30 s, 12 cycles of 98°C for 10 s, 65°C for 30 s, 72°C for 70 s, and the final elongation at 72°C for 5 min. PCR reactions were pooled and cleaned with Sera‐Mag beads (1.5x) before final DNA quantities for each library were taken using Qubit® 2.0 Fluorometer (Thermo Fisher Scientific, VIC, Australia). Six libraries were prepared separately with sequences run on the Illumina HiSeq 4000 at the Novogene (Hong Kong) obtaining 150 bp paired-end reads.

Ancestral lineage proportion

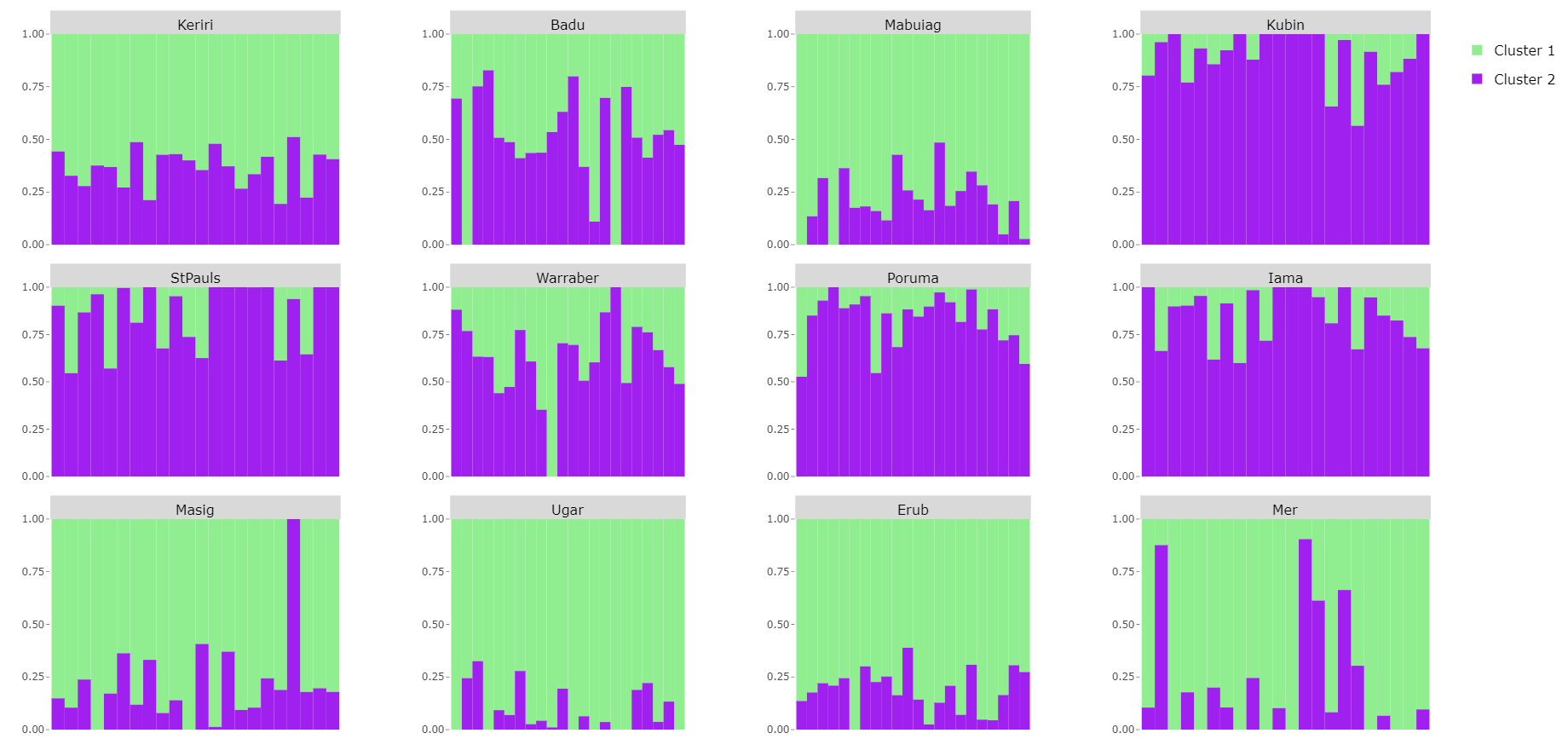

Figure S15: Individual ancestral lineage proportion for each village (n=22) with two ancestral lineages (K = 2) selected.

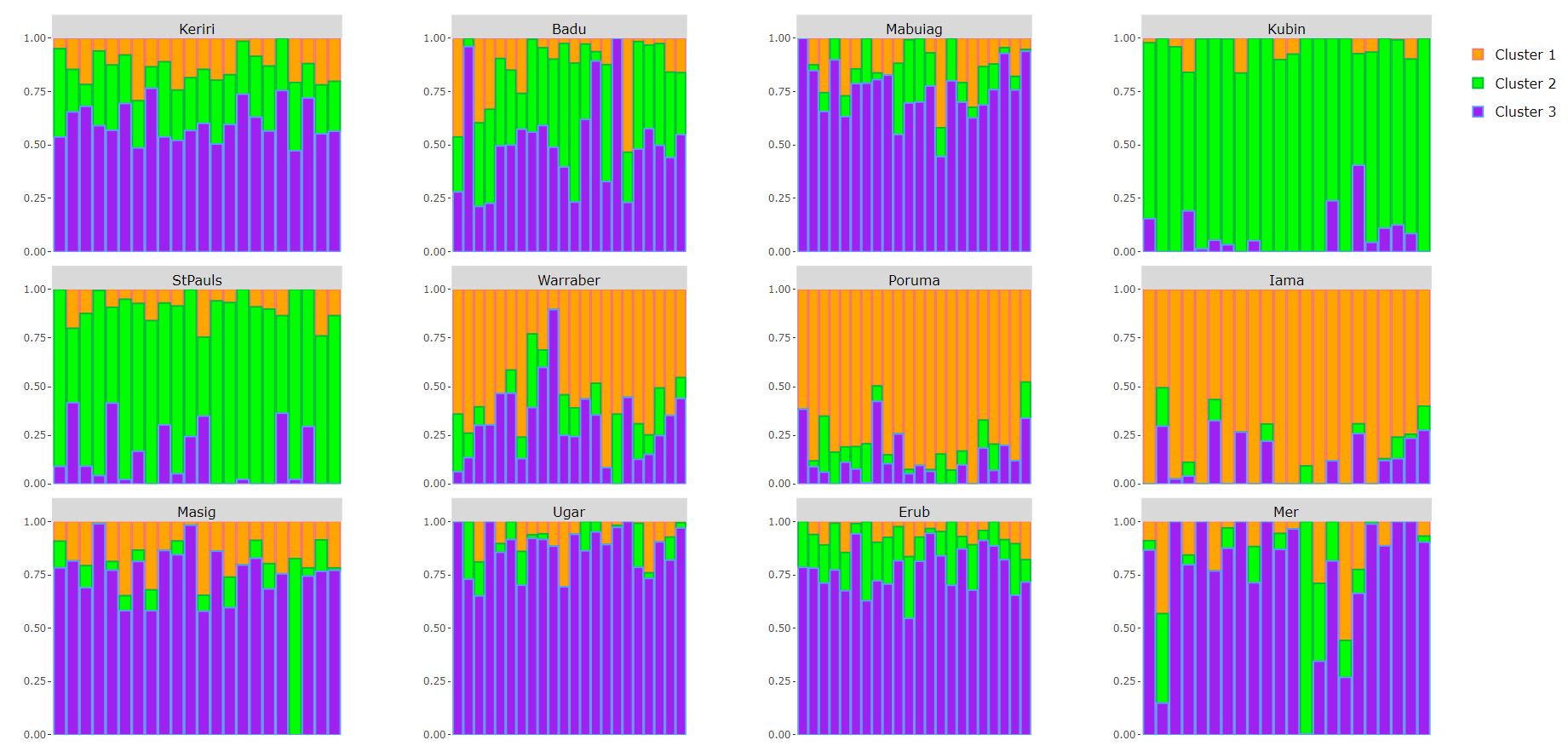

Ancestral lineage proportion

Figure S16: Individual ancestral lineage proportion for each village (n=22) with three ancestral lineages (K = 3) selected.

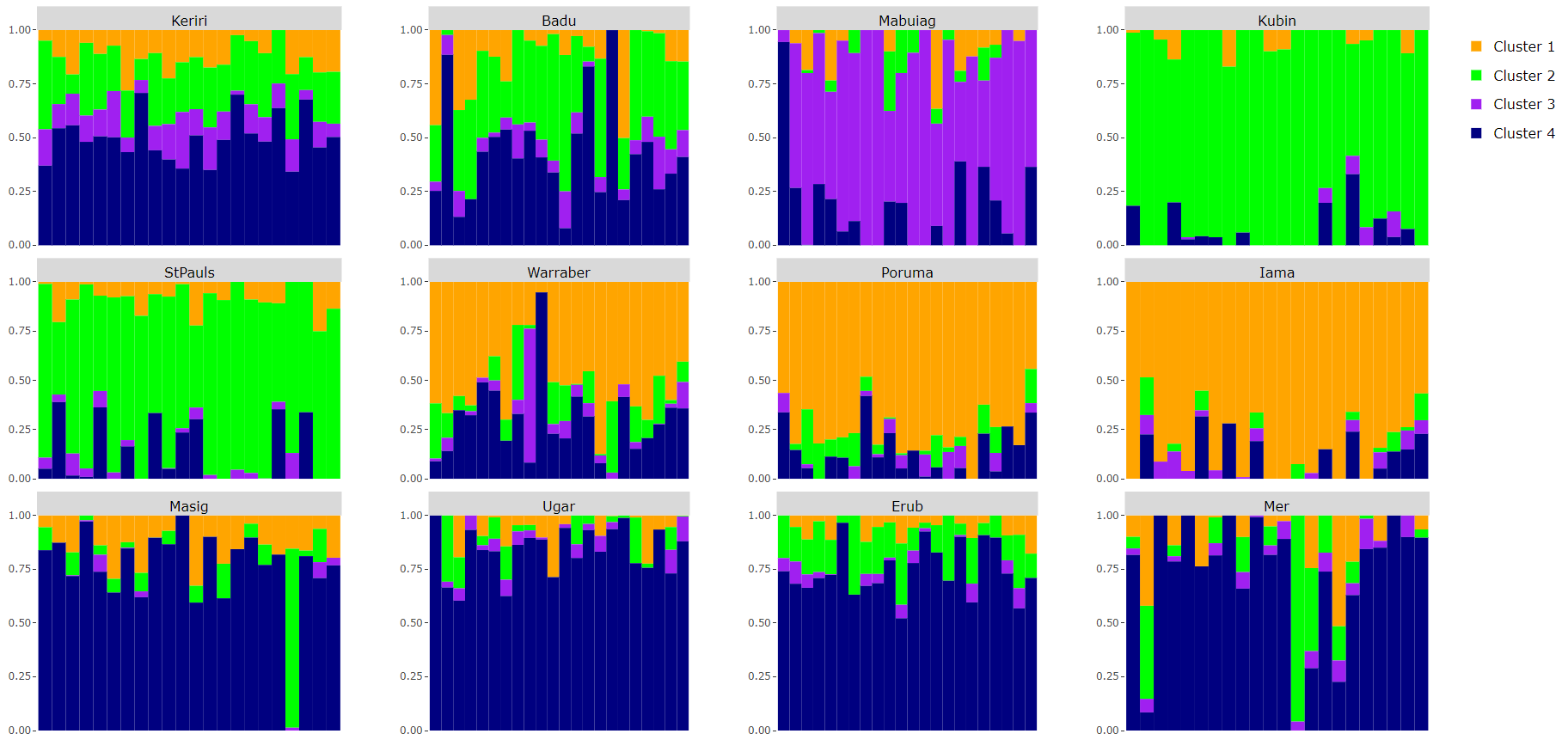

Ancestral lineage proportion

Figure S17: Individual ancestral lineage proportion for each village (n=22) with four ancestral lineages (K = 4) selected.

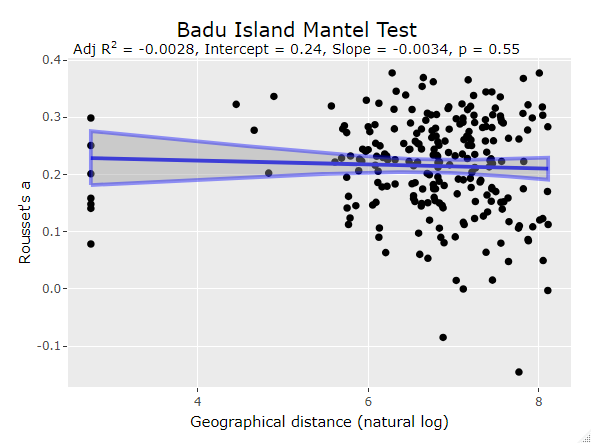

Figure S18: Scatter plot of a mantel test between geographic (natural log) and genetic distance (Rousset’s a) for Aedes albopictus collections on Badu Island. Lines describe a linear regression fit with 95% Confidence Interval shown.

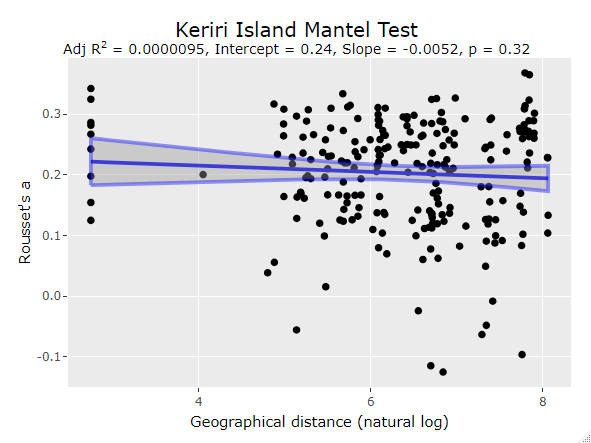

Figure S19: Scatter plot of a mantel test between geographic (natural log) and genetic distance (Rousset’s a) for *Aedes albopictus* collections on Keriri Island. Lines describe a linear regression fit with 95% Confidence Interval shown.

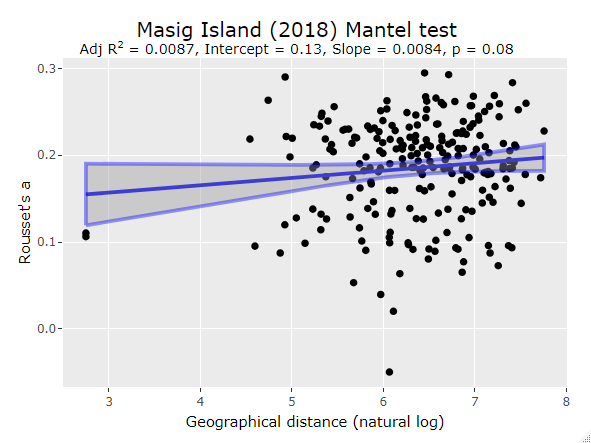

Figure S20: Scatter plot of a mantel test between geographic (natural log) and genetic distance (Rousset’s a) for Aedes albopictus collections on Masig Island in 2018. Lines describe a linear regression fit with 95% Confidence Interval shown.

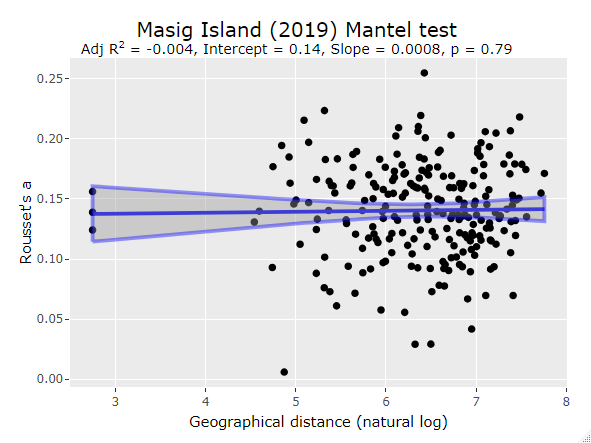

Figure S21: Scatter plot of a mantel test between geographic (natural log) and genetic distance (Rousset’s a) for Aedes albopictus collections on Masig Island in 2019. Lines describe a linear regression fit with 95% Confidence Interval shown.

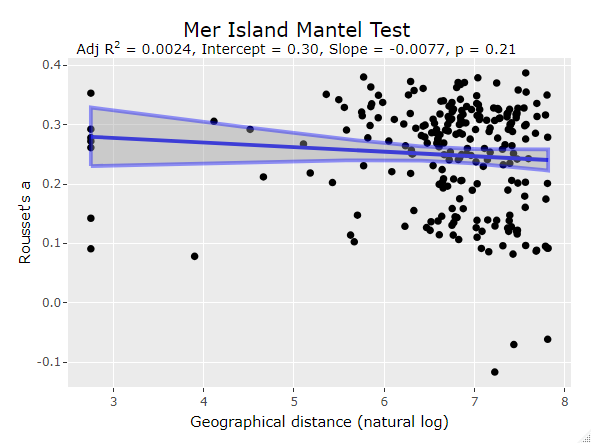

Figure S22: Scatter plot of a mantel test between geographic (natural log) and genetic distance (Rousset’s a) for *Aedes albopictus* collections on Mer Island. Lines describe a linear regression fit with 95% Confidence Interval shown.

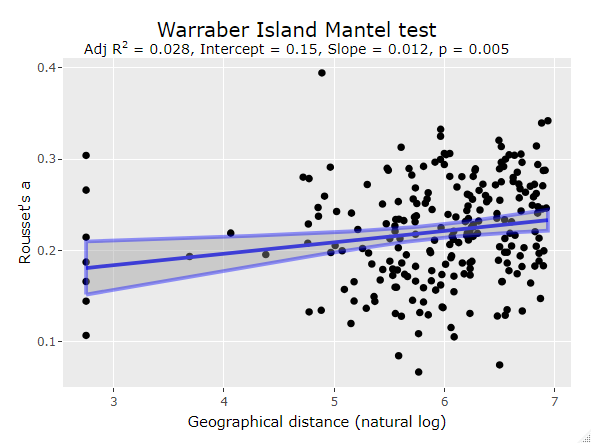

Figure S23: Scatter plot of a mantel test between geographic (natural log) and genetic distance (Rousset’s a) for *Aedes albopictus* collections on Warraber Island. Lines describe a linear regression fit with 95% Confidence Interval shown.

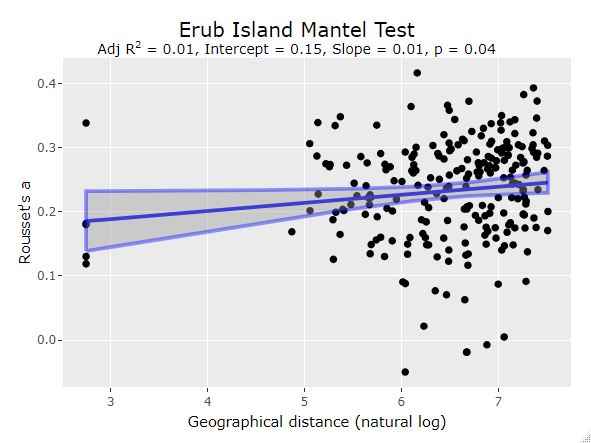

Figure S24: Scatter plot of a mantel test between geographic (natural log) and genetic distance (Rousset’s a) for *Aedes albopictus* collections on Erub Island. Lines describe a linear regression fit with 95% Confidence Interval shown.

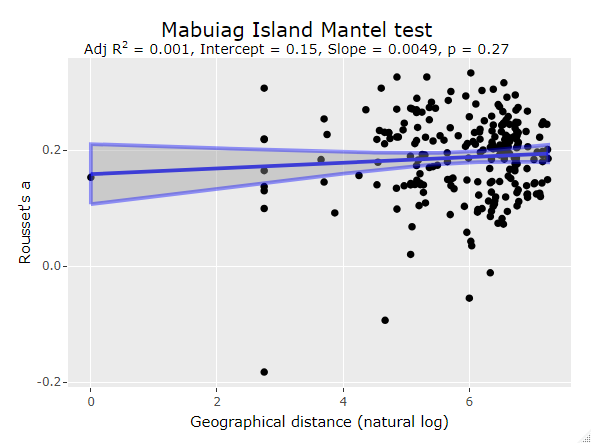

Figure S25: Scatter plot of a mantel test between geographic (natural log) and genetic distance (Rousset’s a) for *Aedes albopictus* collections on Mabuiag Island. Lines describe a linear regression fit with 95% Confidence Interval shown.

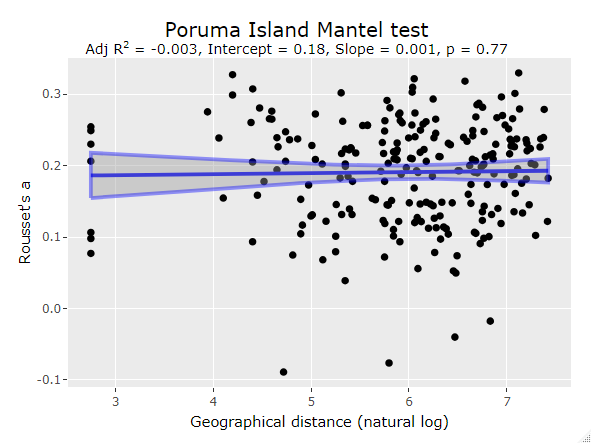

Figure S26: Scatter plot of a mantel test between geographic (natural log) and genetic distance (Rousset’s a) for *Aedes albopictus* collections on Poruma Island. Lines describe a linear regression fit with 95% Confidence Interval shown.

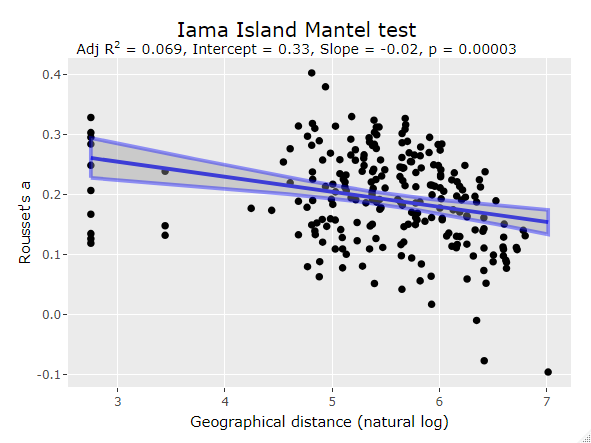

Figure S27: Scatter plot of a mantel test between geographic (natural log) and genetic distance (Rousset’s a) for *Aedes albopictus* collections on Iama Island. Lines describe a linear regression fit with 95% Confidence Interval shown.

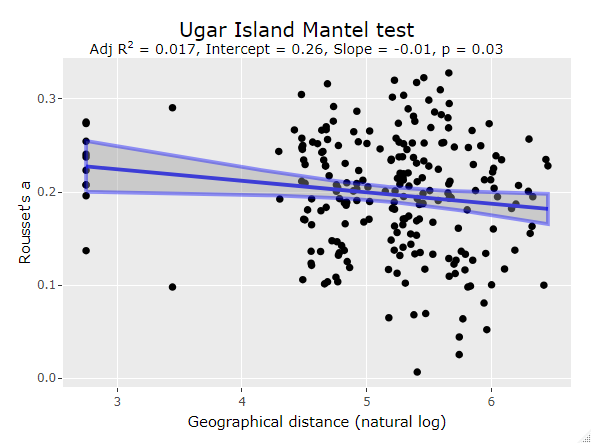

Figure S28*:* Scatter plot of a mantel test between geographic (natural log) and genetic distance (Rousset’s a) for *Aedes albopictus* collections on Ugar Island. Lines describe a linear regression fit with 95% Confidence Interval shown.

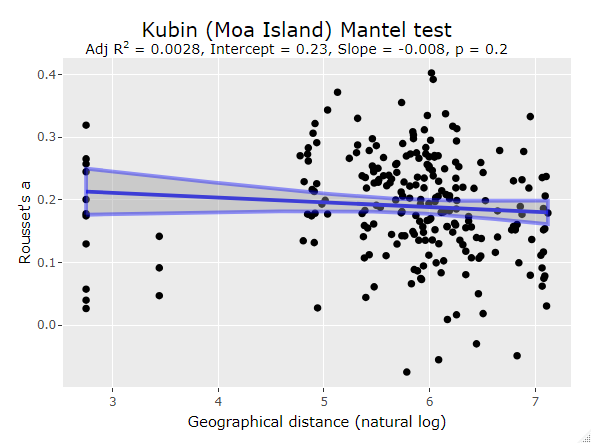

Figure S29: Scatter plot of a mantel test between geographic (natural log) and genetic distance (Rousset’s a) for *Aedes albopictus* collections on Kubin (Moa Island). Lines describe a linear regression fit with 95% Confidence Interval shown.

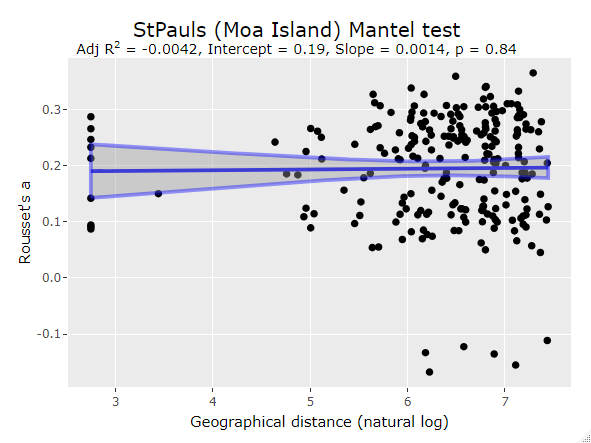

Figure S30: Scatter plot of a mantel test between geographic (natural log) and genetic distance (Rousset’s a) for *Aedes albopictus* collections on St Pauls (Moa Island). Lines describe a linear regression fit with 95% Confidence Interval shown.

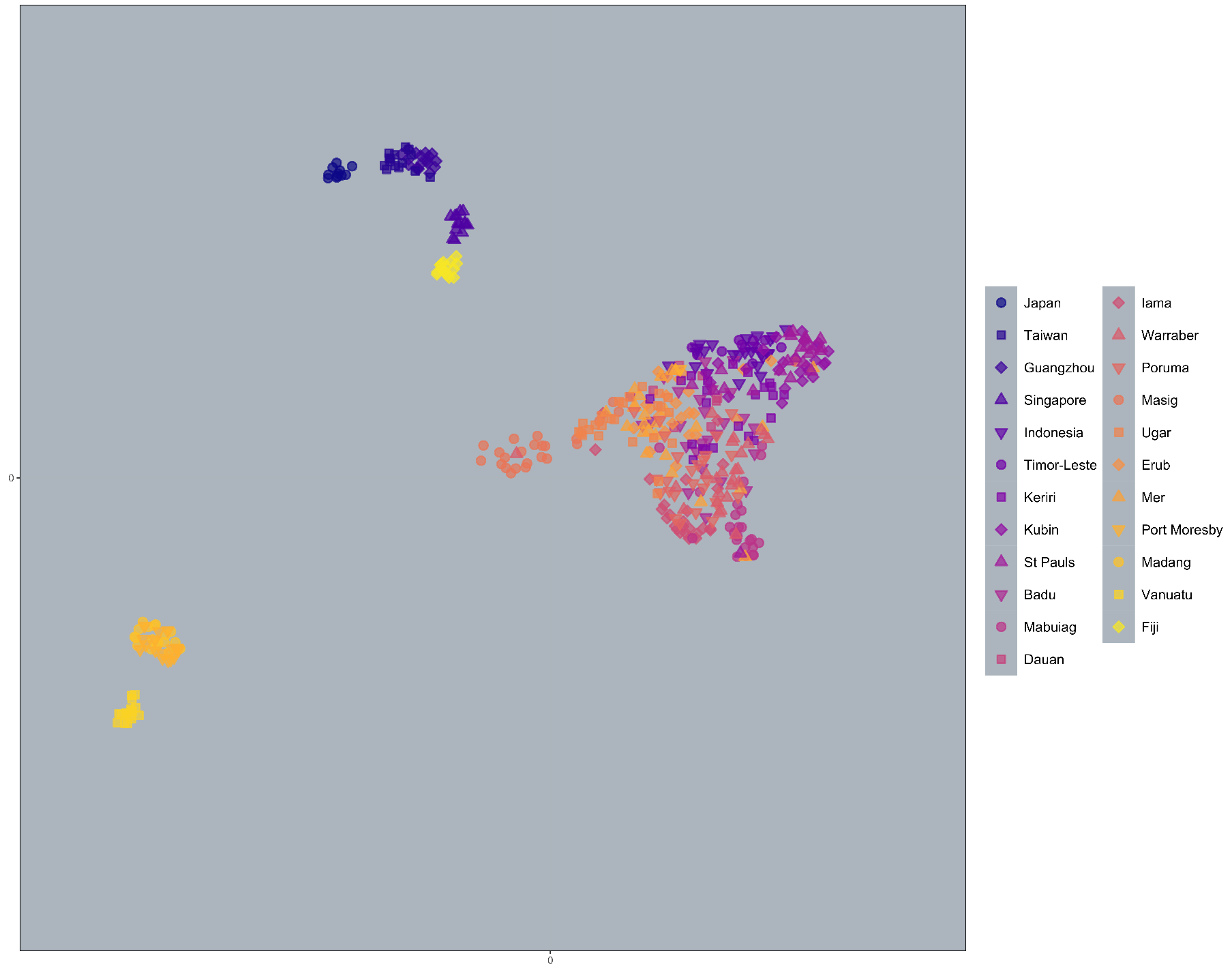

Figure S32: PCA-UMAP of all non-incursive genotypes, using the first 4 PCs, 50 neighbours and a 0.5 minimum distance.

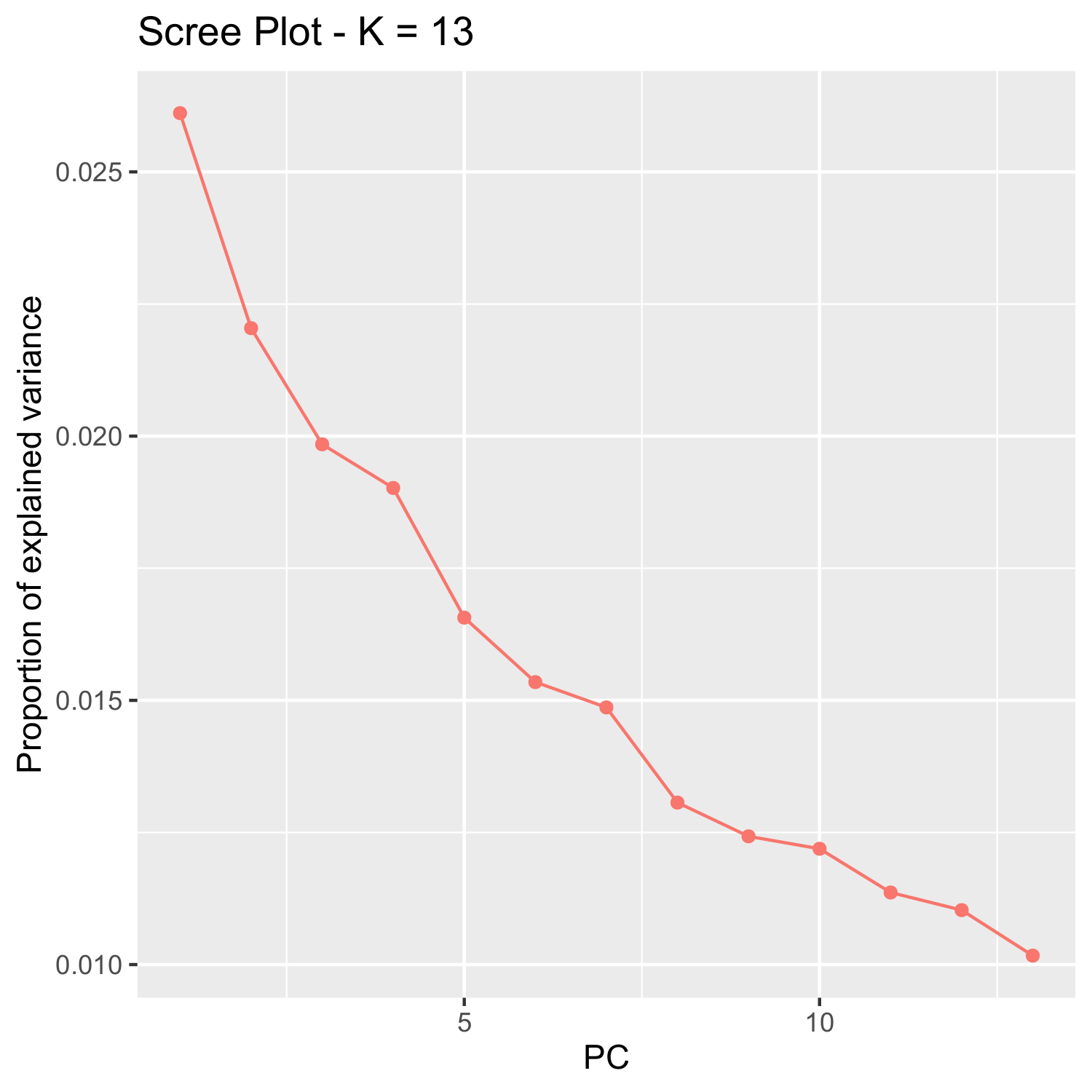

Figure S33: Scree plot from pcadapt.

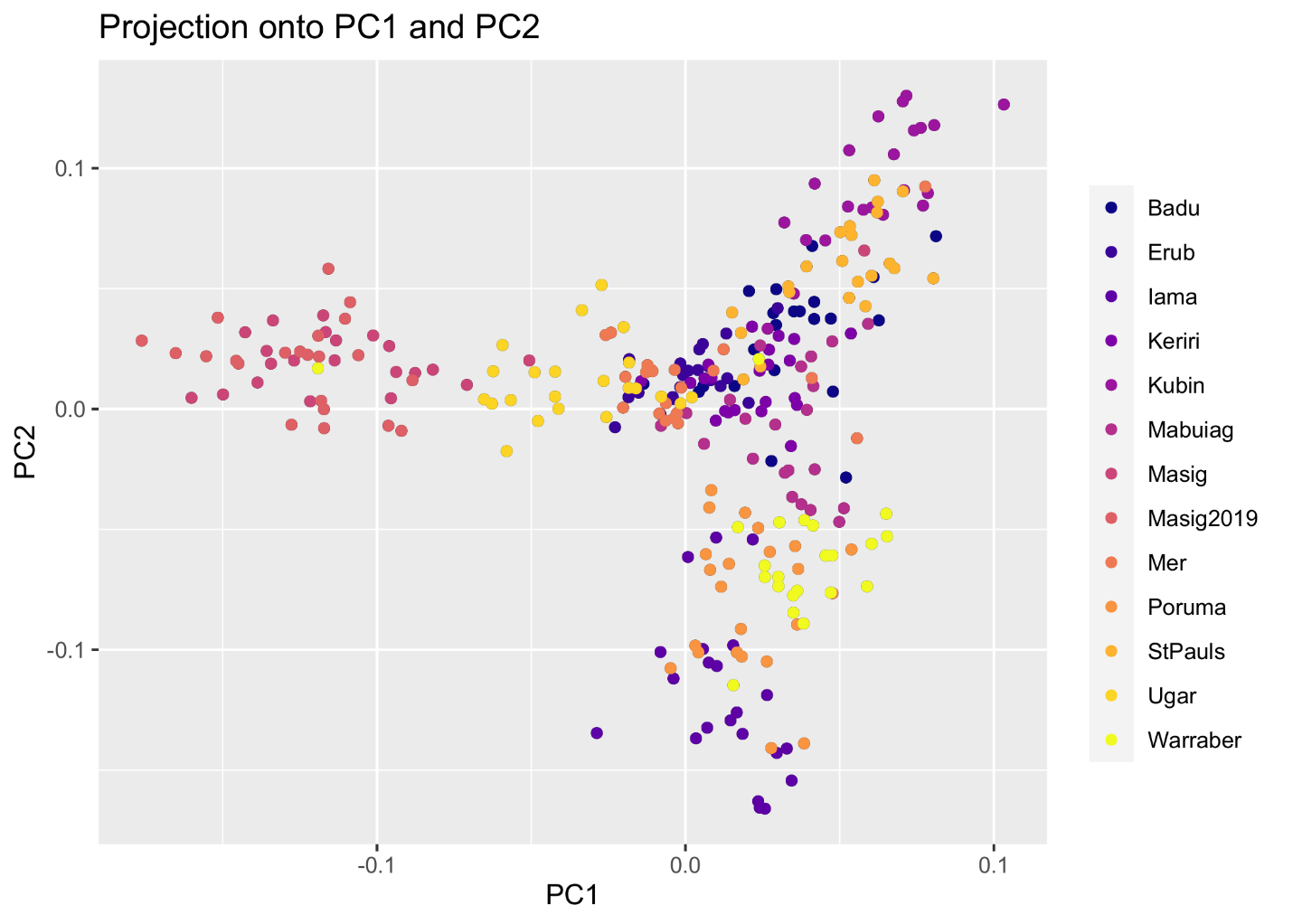

Figure S34: Genome-wide genetic structure of TSI villages, for the first and second PCs.

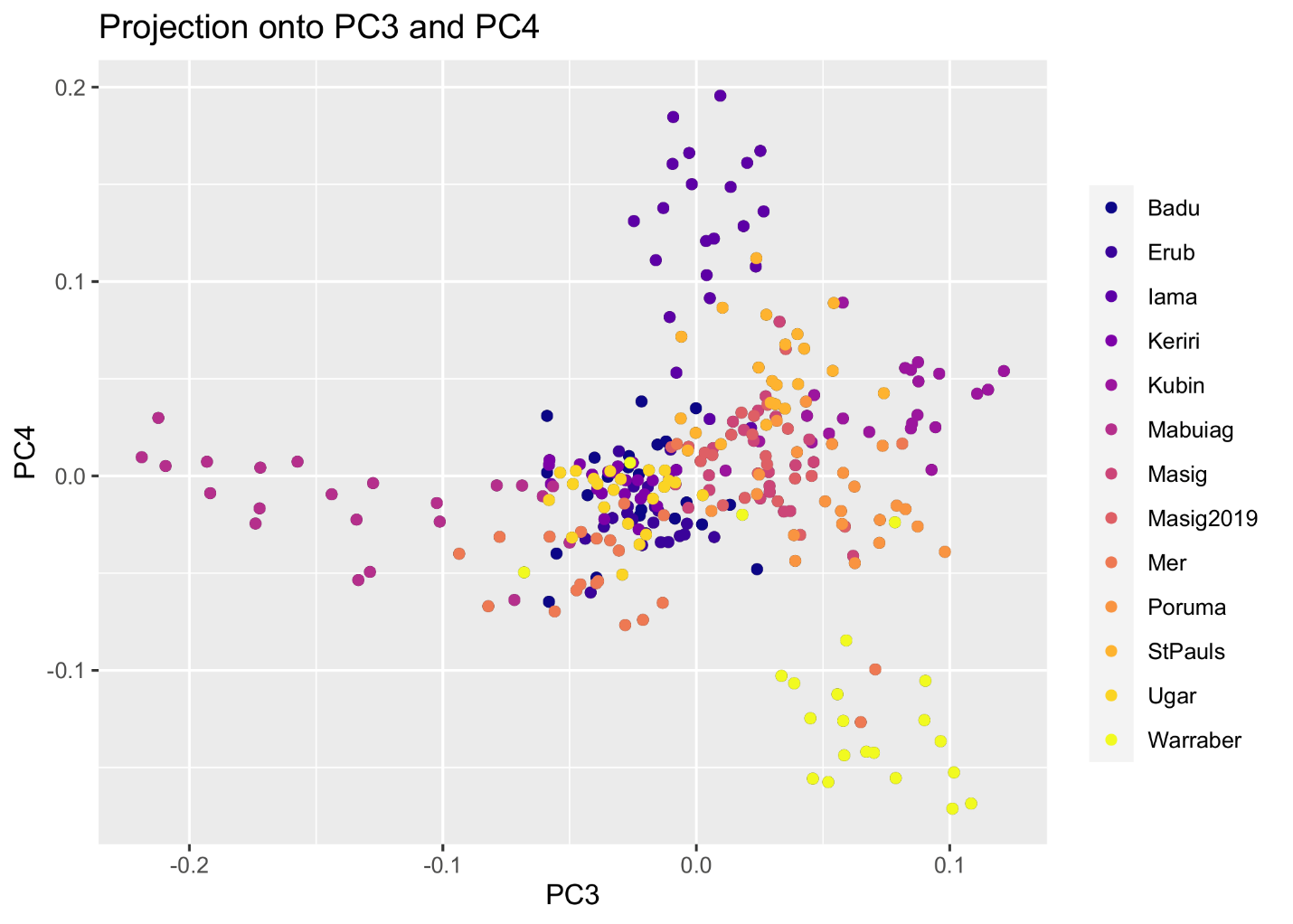

Figure S35: Genome-wide genetic structure of TSI villages, for the third and fourth PCs.

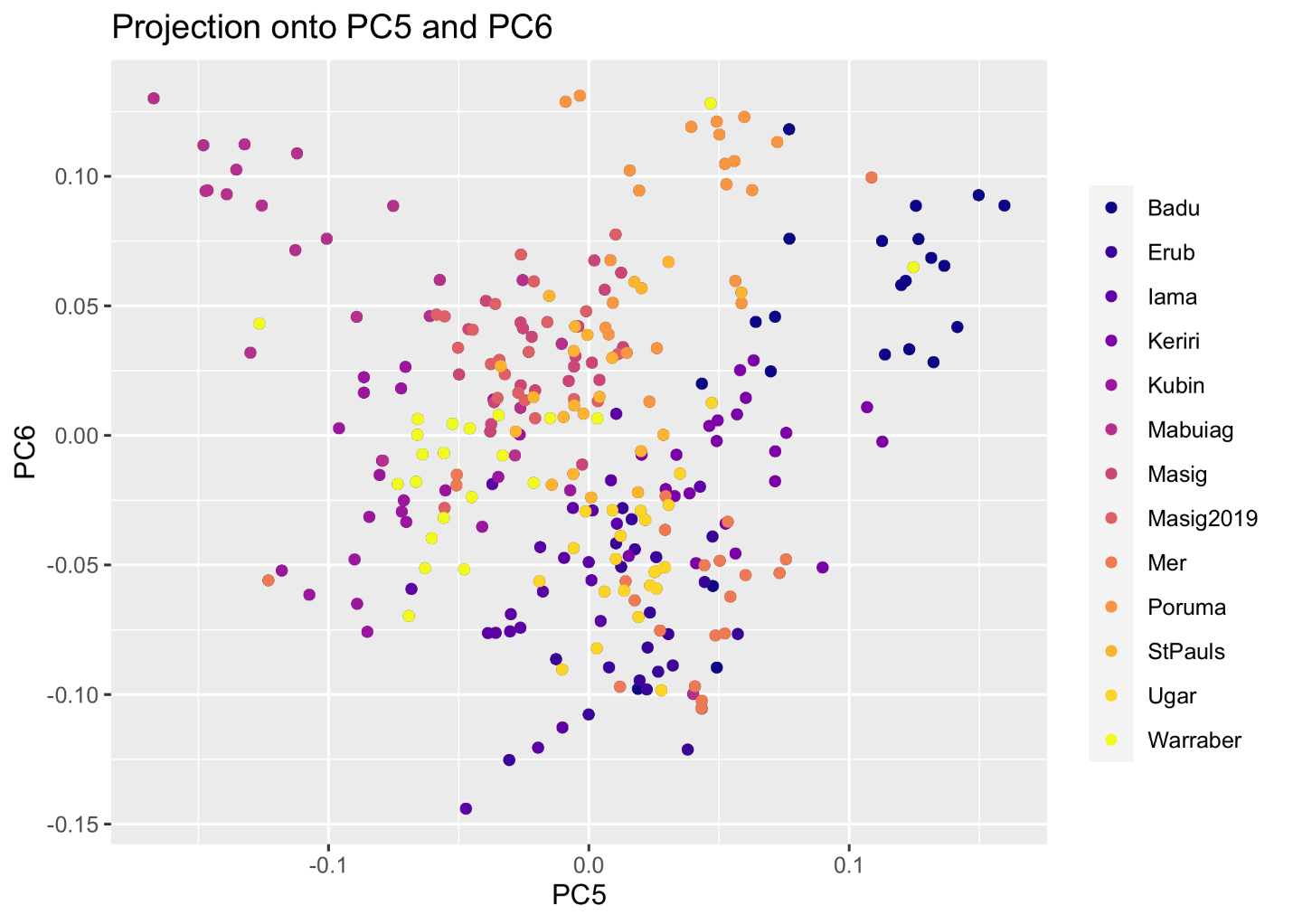

Figure S36: Genome-wide genetic structure of TSI villages, for the fifth and sixth PCs.

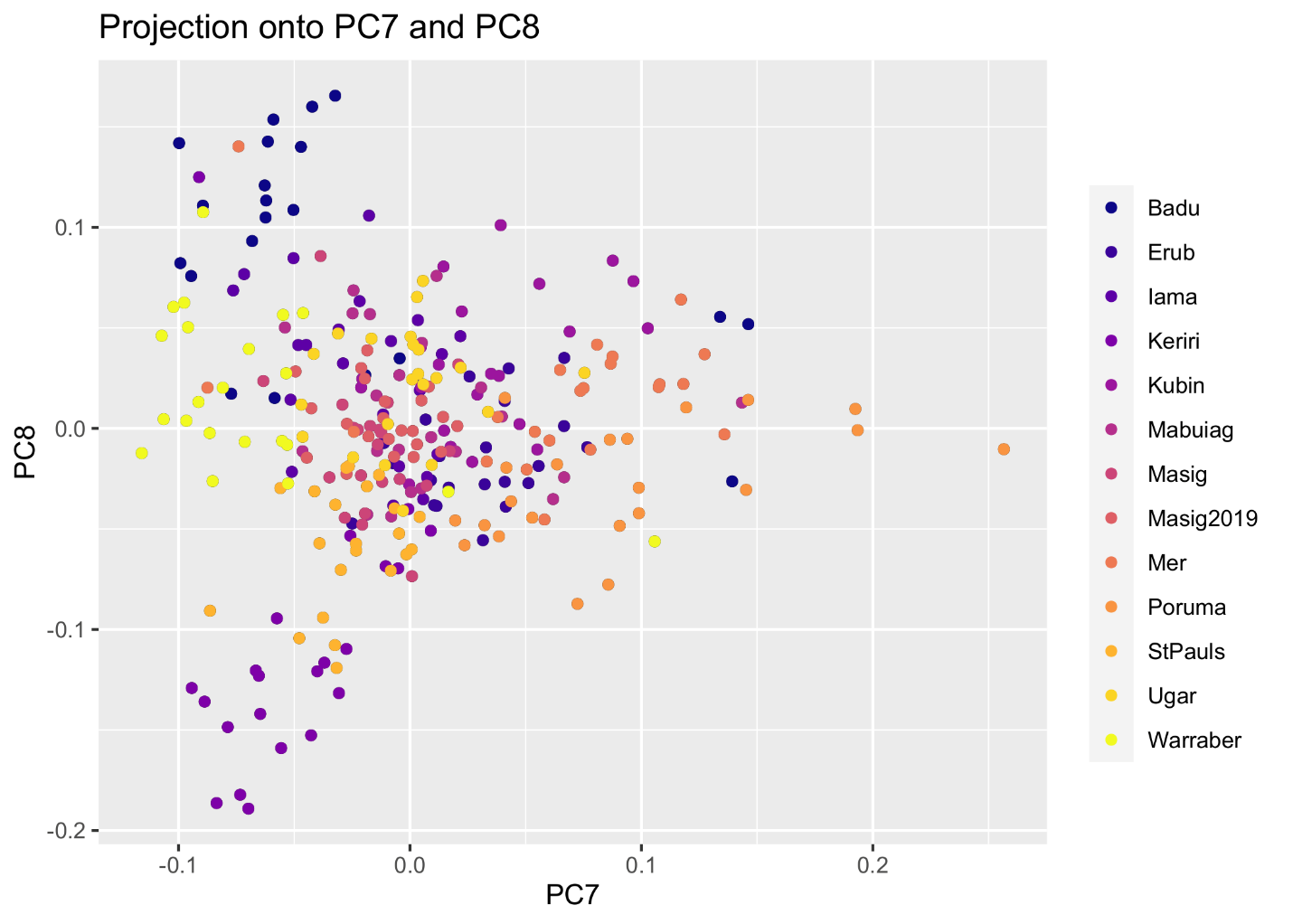

Figure S37: Genome-wide genetic structure of TSI villages, for the seventh and eigth PCs.

Table S1: Initial assignPOP run showing assignment probability to each TSI and non-TSI location, and aggregated assignment probabilities for all TSI and all non-TSI locations.

| Location mosquito | Inc-1 | Inc-2 | Inc-3 | Inc-4 |
| --- | --- | --- | --- | --- |
| **TSI** |  |  |  |  |
| Badu | 0.064 | 0.012 | 0.143 | 0.297 |
| Dauan | 0.032 | 0.062 | 0.048 | 0.038 |
| Erub | 0.036 | 0.039 | 0.044 | 0.033 |
| Iama | 0.015 | 0.039 | 0.025 | 0.047 |
| Keriri | 0.278 | 0.309 | 0.202 | 0.158 |
| Kubin | 0.036 | 0.010 | 0.038 | 0.037 |
| Mabuiag | 0.006 | 0.015 | 0.015 | 0.009 |
| Masig | 0.016 | 0.072 | 0.010 | 0.007 |
| Mer | 0.026 | 0.009 | 0.033 | 0.031 |
| Poruma | 0.012 | 0.040 | 0.017 | 0.009 |
| St Pauls | 0.049 | 0.020 | 0.079 | 0.060 |
| Ugar | 0.300 | 0.305 | 0.192 | 0.153 |
| Warraber | 0.059 | 0.023 | 0.048 | 0.059 |
| **Non-TSI** |  |  |  |  |
| Bali | 0.003 | 0.007 | 0.006 | 0.003 |
| Bandung | 0.003 | 0.003 | 0.006 | 0.003 |
| Fiji | 0.004 | 0.002 | 0.007 | 0.005 |
| Guangzhou | 0.006 | 0.003 | 0.007 | 0.006 |
| Jakarta | 0.003 | 0.005 | 0.012 | 0.004 |
| Japan | 0.004 | 0.002 | 0.005 | 0.004 |
| Madang | 0.005 | 0.002 | 0.006 | 0.004 |
| Port Moresby | 0.026 | 0.011 | 0.034 | 0.019 |
| Singapore | 0.004 | 0.002 | 0.007 | 0.004 |
| Taiwan | 0.007 | 0.003 | 0.007 | 0.006 |
| Timor-Leste | 0.003 | 0.003 | 0.004 | 0.002 |
| Vanuatu | 0.003 | 0.002 | 0.005 | 0.003 |
| **TSI (all)** | **0.928** | **0.956** | **0.893** | **0.937** |
| **Non-TSI (all)** | **0.072** | **0.044** | **0.107** | **0.063** |

Table S2: Pairwise FST estimates among TSI villages.

|  | Badu | Erub | Iama | Keriri | Kubin | Mabuiag | Masig | Masig2019 | Mer | Poruma | StPauls | Ugar | Warraber |
| --- | --- | --- | --- | --- | --- | --- | --- | --- | --- | --- | --- | --- | --- |
| Badu |  | 0.026 | 0.034 | 0.025 | 0.030 | 0.029 | 0.028 | 0.031 | 0.022 | 0.029 | 0.027 | 0.027 | 0.028 |
| Erub |  |  | 0.033 | 0.026 | 0.031 | 0.030 | 0.027 | 0.029 | 0.022 | 0.030 | 0.029 | 0.026 | 0.029 |
| Iama |  |  |  | 0.034 | 0.040 | 0.036 | 0.034 | 0.036 | 0.032 | 0.032 | 0.035 | 0.034 | 0.035 |
| Keriri |  |  |  |  | 0.032 | 0.030 | 0.029 | 0.031 | 0.025 | 0.030 | 0.028 | 0.028 | 0.029 |
| Kubin |  |  |  |  |  | 0.036 | 0.033 | 0.036 | 0.030 | 0.035 | 0.028 | 0.033 | 0.033 |
| Mabuiag |  |  |  |  |  |  | 0.032 | 0.034 | 0.027 | 0.033 | 0.032 | 0.030 | 0.032 |
| Masig |  |  |  |  |  |  |  | 0.014 | 0.025 | 0.030 | 0.029 | 0.025 | 0.029 |
| Masig2019 |  |  |  |  |  |  |  |  | 0.027 | 0.032 | 0.033 | 0.027 | 0.032 |
| Mer |  |  |  |  |  |  |  |  |  | 0.028 | 0.028 | 0.025 | 0.025 |
| Poruma |  |  |  |  |  |  |  |  |  |  | 0.032 | 0.031 | 0.028 |
| StPauls |  |  |  |  |  |  |  |  |  |  |  | 0.031 | 0.032 |
| Ugar |  |  |  |  |  |  |  |  |  |  |  |  | 0.030 |
